# A One Health approach metagenomic study on the antimicrobial resistance traits of canine saliva

**DOI:** 10.1101/2024.01.17.576087

**Authors:** Adrienn Gréta Tóth, Darinka Lilla Tóth, Laura Remport, Imre Tóth, Tibor Németh, Attila Dubecz, Árpád V. Patai, László Makrai, Norbert Solymosi

## Abstract

According to the One Health concept, physical proximity among pets and their owners facilitates the spread of various bacteria. Interspecies bacterial transfer media include canine saliva that can be transmitted by licking and biting. Antimicrobial resistance genes (ARGs) are the natural constituents of the bacterial genome. However, human activity can increase the prominence of ARGs. To assess factors that may affect the resistome of the canine oral cavity, the shotgun metagenomic sequencing dataset of 1830 canine saliva samples was bioinformatically analyzed and supplemented with survey results of the physical and behavioral traits of the dogs. Bacteriome and resistome associated with the canine saliva samples were described throughout the analysis. Moreover, the subset of higher public health ARGs and ESKAPE pathogen-related (Enterococcus faecium, Staphylococcus aureus, Klebsiella pneumoniae, Acinetobacter baumannii, Pseudomonas aeruginosa, and Enterobacter species) higher public health ARGs were also collected. Further on, the set and subsets of ARGs were studied considering the surveyed traits of the sampled dogs. Overall, 318 ARG types reached sufficiently high detection rates. These ARGs can affect 31 antibiotic drug classes by various resistance mechanisms. ARGs against tetracyclines and cephalosporins appeared in the highest number of samples. However, surprisingly, another CIA group, peptides were represented by ARGs in the third-highest number of samples. Considering further ARG appearance rates in the samples, Critically Important Antimicrobials (CIAs, WHO), such as aminoglycosides, fluoroquinolones, or macrolides, were among the most frequently affected drug classes by higher public health risk ARGs and ESKAPE pathogen related higher public health risk ARGs. Bacteria in the saliva of white and diluted (merle, gray) color dogs and dogs characterized with decreased activity and decreased aggression more often harbored ARGs. Reduced playfulness could have been specifically associated with higher public health risk ARG presence. Even though the oral microbiome of the owners is unknown, One Health and public health implications of the close human-pet bonds and factors potentially underlying the rise in salivary ARG numbers should be considered, mostly in the light of the presence of ARGs affecting critically important drugs for human medicine.

## Introduction

The interconnection of human, animal and environmental habitats contributes the evolution of one of the most concerning global health issues, antimicrobial resistance (AMR). According to the One Health concept, the spread of resistant organisms or their AMR determinants can occur among human-associated, animal-associated and environmental microbiomes in certain conditions (e.g. administration of antibiotics). Transport media involved in local AMR transmission pathways among physically connected habitats can be screened for their role in the growing appearance rates of AMR. Such a pathway is among humans and their companion animals.

After a decade-long rise in the number of companion animals in households, more than 50% of the U.S. population held a dog in 2020.^1–3^ Moreover, a change in the quality of human-pet bonds is also notable. According to the survey of the American Veterinary Medical Association, 70% of pet owners consider their pets as family members, 17% as companions and only 3% as property.^4^ This mindset enhances physical proximity (e.g. sleeping with the owner, licking the face of the owner etc.) among the pets and their owners.^5^ Companion dogs, that are often treated as family members, undergo regular veterinary healthcare services. Such services can include treatments with antibiotics. A study from 2016 states that 1 in 4 dogs (25.2%, 95% CI: 25.1–25.3%) from the United Kingdom were treated with antibiotics in a two-year period.^6^ In addition to the growth in pet dog numbers and the closer pet-owner co-existences, dog bite cases are also becoming more frequent.^7–11^ Interestingly, only 2 out 5 dog bites are associated with strays, while the majority of these cases are related to family dogs.^12^

Physical contact by close dog-owner co-existence and by dog bite wounds serve as a platform for interspecies bacterial and AMR determinant transfer. Moreover, this platform may provide opportunity for the interspecies transfer of ESKAPE pathogens identified by the WHO (*Enterococcus faecium*, *Staphylococcus aureus*, *Klebsiella pneumoniae*, *Acinetobacter baumannii*, *Pseudomonas aeruginosa*, and *Enterobacter* spp.) that are antibiotic-resistant "priority pathogens" posing the greatest threat to human health or even higher public health risk ARGs^13^. To categorize ARGs according to their public health risk, Zhang and colleagues^13^ used four indicators: higher human accessibility, mobility, pathogenicity, and clinical availability.

Our aim was to obtain information on the associations of three antimicrobial resistance gene (ARG) sets (all detected ARGs, higher public health risk ARGs^13^, ESKAPE pathogen-related higher public health risk ARGs) in canine saliva and certain easily observable canine traits (e.g. sterilization status, breed, head shape, fur color, behavioural properties). For this purpose, canine shotgun metagenonomic data from the study of Morrill and colleagues^14^ were used. The utilised genomic dataset was linked with answers from forms of canine characteristics and a survey of 118, mostly behavior-related questions filled by the owners of the sampled dogs. ARG abundances were analysed based on phenotypical groups formed based on these answers.

## Results

Results obtained from the bioinformatic analysis of the 1830 canine salivary samples’ metagenome sequencing data were articlulated as follows. After the presentation of the most frequently detected bacterial taxa (bacteriome) associated with ARGs and the set of identified ARGs (resistome), physical and behavioural traits potentially related to the AMR determinants in the canine salivary bacteriome are detailed. Only results with potential biomedical significance are presented in the following sections. Further results are represented among the supplementary materials.

### Bacteriome

By the taxonomic classification, 126 genera could have been identified and associated with ARGs from 1682 samples. ARG associated bacterial genera that appeared in at least 2% of the samples were *Porphyromonas* (33.1%), *Pasteurella* (31.4%), *Frederiksenia* (23.2%), *Pseudomonas* (23.0), *Conchiformibius* (21.7%), *Bacteroides* (10%), *Glaesserella* (9.5%), *Wielerella* (8.7%), *Riemerella* (8.7%), *Escherichia* (8.6%), *Prevotella* (7.3%), *Capnocytophaga* (6.8%), *Klebsiella* (5.2%), *Alistipes* (4.7%), *Lelliottia* (4.5%), *Mannheimia* (4.0%), *Histophilus* (3.9%), *Streptococcus* (3.8%), *Serratia* (3.7%), *Haemophilus* (3.1%), *Enterobacter* (2.6%), *Clostridioides* (2.4%), *Parabacteroides* (2.4%). The number of samples in which these bacterial genera appeared are demostrated on Figure 1.

**Figure 1.**
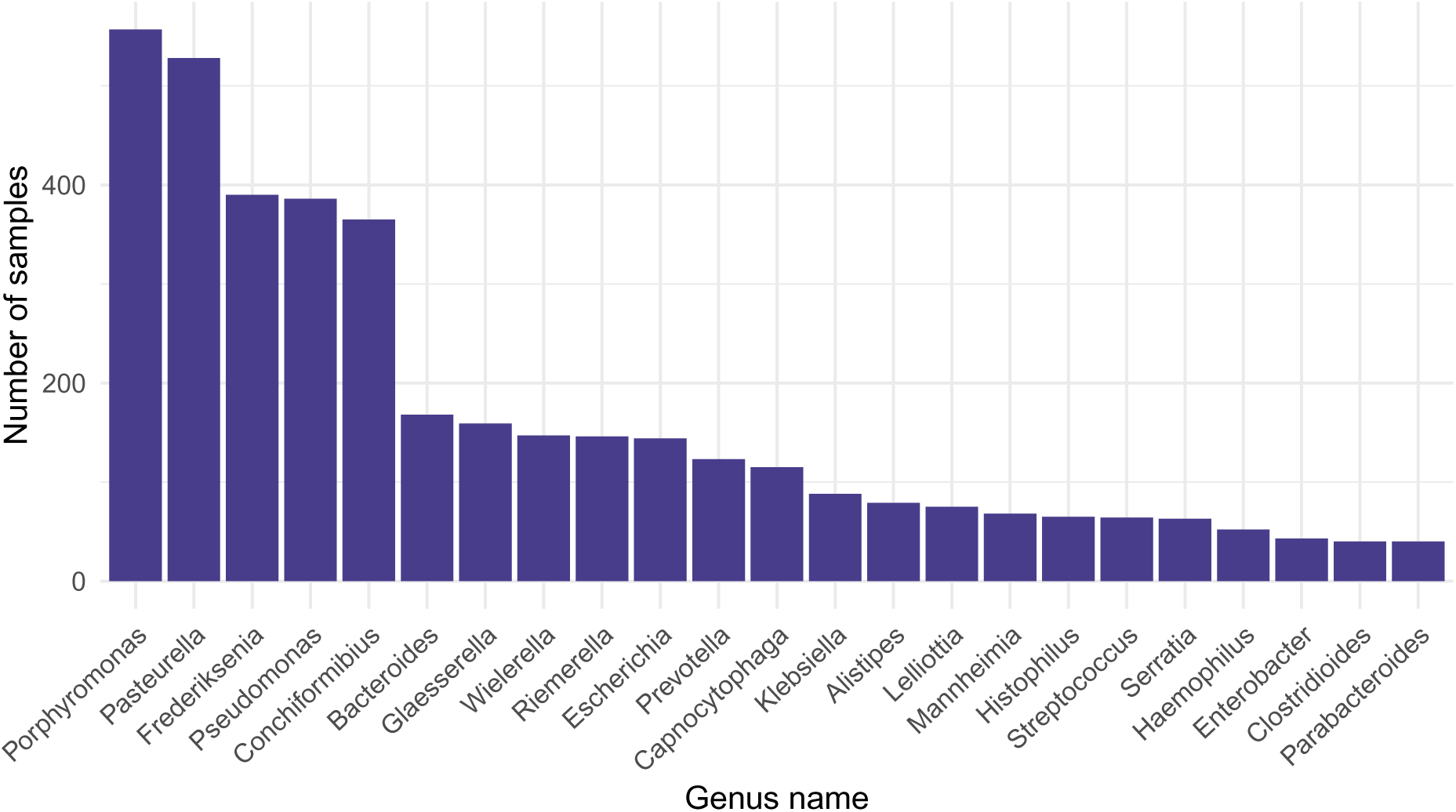
Bacterial genera associated with AMR determinants that appeared in at least 2% of all 1682 samples, with the number of samples in which they were detected.

### Resistome

Setting the 90% length and 90% base identitity tresholds by the ARG identification, 1682 samples could have been associated with ARGs in general, although only 1679 of these samples contained genes that can have antimicrobial effects. Three genes, namely *CRP, nalC* and *nalD* that were identified in the samples based on the CARD database are related to AMR. However, these genes are ARG regulators or repressors and not ARGs. Thus, *CRP, nalC* and *nalD* were excluded from further analysis steps that were targeting the ARG content. The highest number of ARG types (65) was found in BioSample SAMN16779058 and SAMN16778891 (57), followed by SAMN16778736 and SAMN16779288 (55). The mean of ARG hits per sample reached 4.3, and the median was 3. 318 ARG types met the criteria of having 90% coverage and 90% sequential identity. The most common ARGs were *Escherichia coli EF-Tu* mutants conferring resistance to Pulvomycin, *pgpB* and *CfxA2* genes, detected in 1002, 953 and 609 samples, respectively. ARG hits of higher public health significance were detected in 1033 samples. The most public health risk hits were discovered in Biosample SAMN16779288 (35), SAMN16779058 (30) and SAMN16778891 parallel with SAMN16777653 (23). The average number of higher public health risk ARG hits per sample was 2.3, while the median was 1. Of the 65 detected ARGs of higher public health risk, the most common were *CfxA2*, *ErmF* and *APH(6)-Id*, appearing in 609, 202, 153 samples, respectively. ESKAPE pathogen-related ARG hits of higher public health risk were identified in 63 samples. Biosample SAMN16778736 contained the most hits of this type (15), followed by SAMN16777571 (14) and SAMN16778717 (13). The mean of ESKAPE pathogen-related higher public health risk ARG hits per sample amounted to 3.3, and the median to 2. All in all, 13 types of ARG hits of public health risk were detected in ESKAPE pathogens. Of these, the most common were *OXA-2* in 18 samples, *APH(6)-Id* in 17 samples and *sul1* 15 samples. 20.4% of all ARG types were higher public health risk ARGs. Examining ESKAPE pathogen-related higher public health risk ARG types, proportions amounted to 4% considering all ARG hits and 20% of higher public health risk ARG types. ARGs appearing in more than 10 samples together with higher public health risk ARGs and higher public health risk ARGs associated to ESKAPE pathogens that appeared in at least 5 samples are demonstrated on Figure 2. The full list of ARGs and the numbers of samples which they have appeared at can be seen at Supplementary table 5.

**Figure 2.**
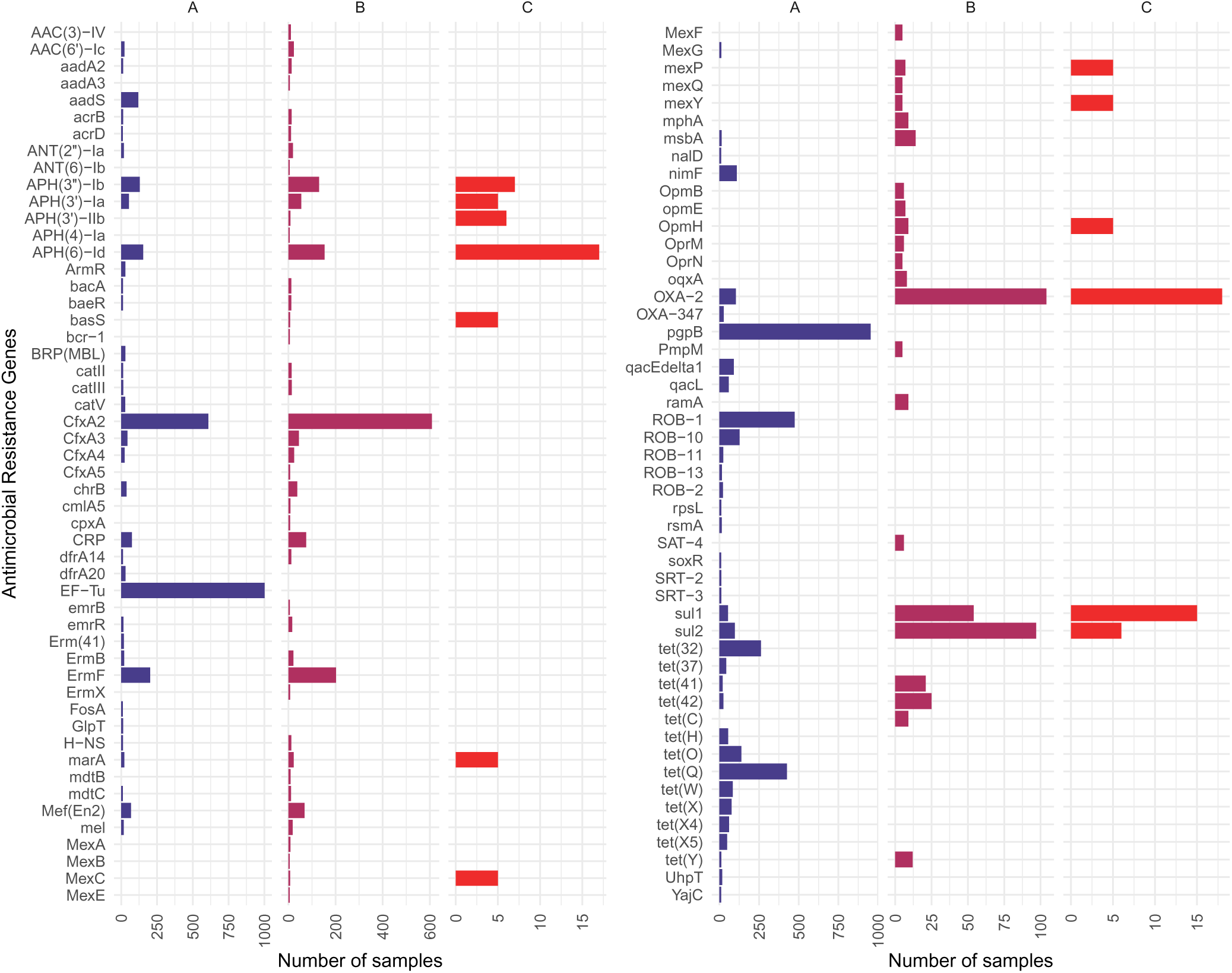
ARGs detected in more than 10 samples (A), higher public health risk ARGs detected in at least 5 samples (B) and higher public health risk ARGs deriving from ESKAPE pathogens detected in at least 5 samples (C). *EF-Tu* is the abbreviation for *Escherichia coli EF-Tu* mutants conferring resistance to Pulvomycin, *UhpT* for *E. coli UhpT* with mutation conferring resistance to fosfomycin, *GlpT* for *E. coli GlpT* with mutation conferring resistance to fosfomycin, *soxR* for *Pseudomonas aeruginosa soxR*, and *rpsL* for *Mycobacterium tuberculosis rpsL* mutations conferring resistance to streptomycin.

Drug classes that could potentially be affected by the presence of the detected ARGs are presented on Figure 3 considering all ARGs, higher public health risk ARGs and higher public health risk ARGs in ESKAPE pathogens. ARGs detected in the samples can potentially affect 31 antibiotic classes, including aminocoumarins, aminoglycosides, bicyclomycins, carbapenems, cephalosporins, cephamycin, diaminopyrimidines, disinfecting agents and antiseptics, elfamycins, fluoroquinolones, fosfomycin, glycopeptides, glycylcyclines, lincosamides, macrolides, monobactams, nitrofurans, nitroimidazoles, nucleosides, oxazolidinones, penams, penems, peptides, phenicols, pleuromutilins, rifamycins, streptogramin A, streptogramins, streptogramin B, sulfonamides and tetracyclines. The most commonly affected antimicrobial groups in descending order were tetracyclines (1650 samples), cephalosporins (1125 samples), peptides (1105 samples) and penams (1095 samples) considering all the detected ARGs. In case of higher public health risk ARGs, cephamycin resistance genes appeared the most (754 samples), followed by macrolides (529 samples) and amynoglycosides (508 samples). By higher public health risk ARGs appearing in ESKAPE pathogens amynoglycoside ARGs (61 samples) were the most common, followed by fluoroquinolones (59 samples), phenicols (54 samples), penams (53 samples) and tetracyclines (53 samples). Considering all three groups (all ARGs, higher public health risk ARGs, higher public health risk ARGs in ESKAPE pathogens) pleuromutilin and oxazolidinone resistance appeared in the lowest number of samples. The exact sample numbers by antibiotic groups can be seen in Supplementary table 6.

**Figure 3.**
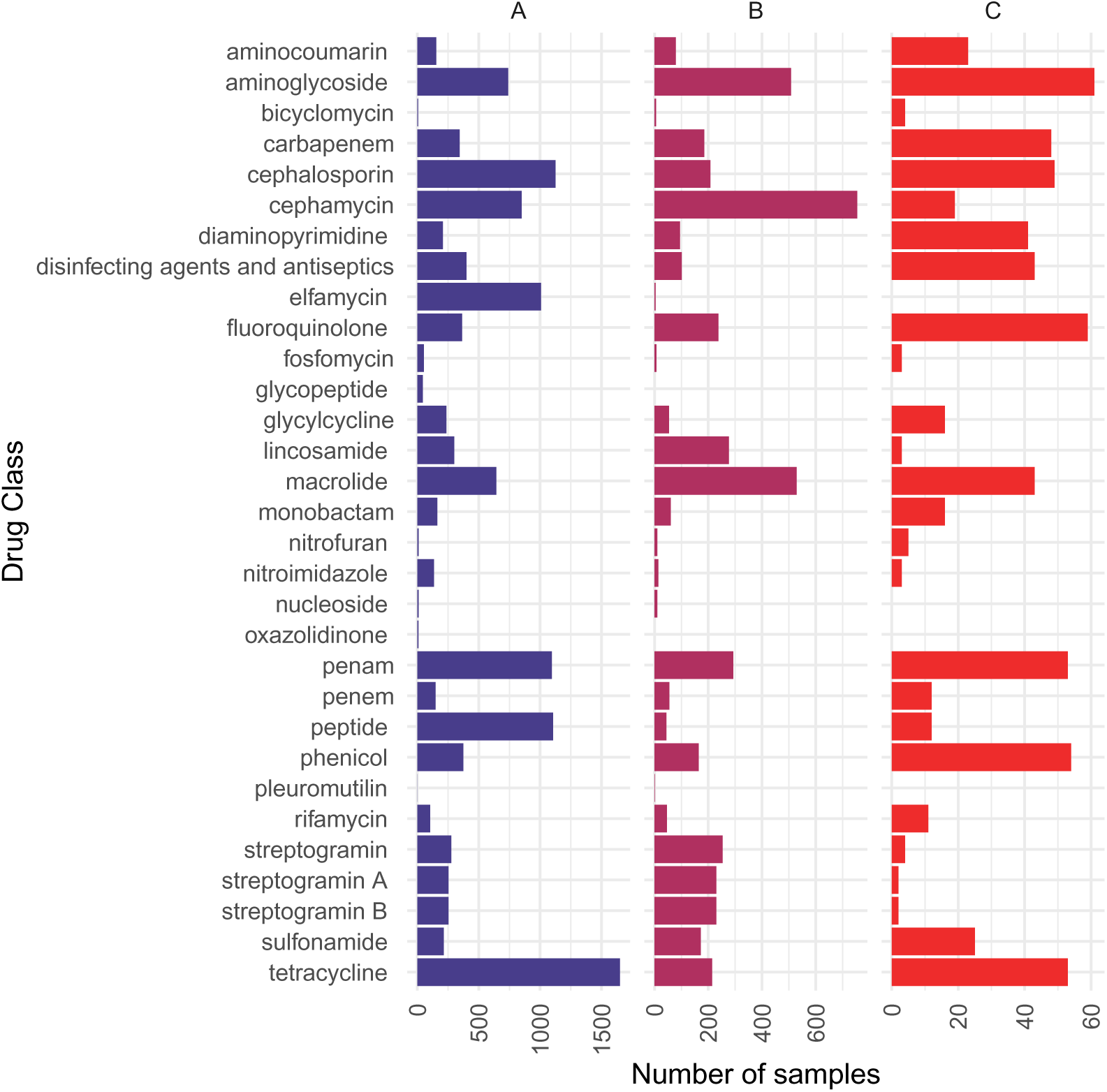
Antibiotic groups against which ARGs were detected in any metagenomic samples (A), antibiotic groups against which higher public health risk ARGs were detected in any metagenomic samples (B) and antibiotic groups against which higher public health risk ARGs appeared in ESKAPE pathogens (C). The number of samples in which ARGs against the presented antibiotic groups were detected is presented on the horizontal axis. Antibiotic compounds affected by multidrug resistance are displayed separately.

The proportion of resistance mechanisms was calculated based on the diversity of all ARGs, higher public health risk ARGs and higher public health risk ARGs in ESKAPE pathogens. The results are demonstrated on Figure 4. The dominant mechanism of identified ARGs was antibiotic inactivation (2669 occasions), antibiotic target alteration (2403 occasions) and antibiotic efflux (973 occasions). In case of higher public health risk ARGs, the top AMR mechanisms were antibiotic inactivation (1304 occasions), antibiotic efflux (533 occasions) and antibiotic target alteration (292 occasions). Higher public health risk ARGs identified in ESKAPE pathogens were in most cases associated with antibiotic efflux (110 occasions), antibiotic inactivation and antibiotic target replacement (25 occasions).

**Figure 4.**
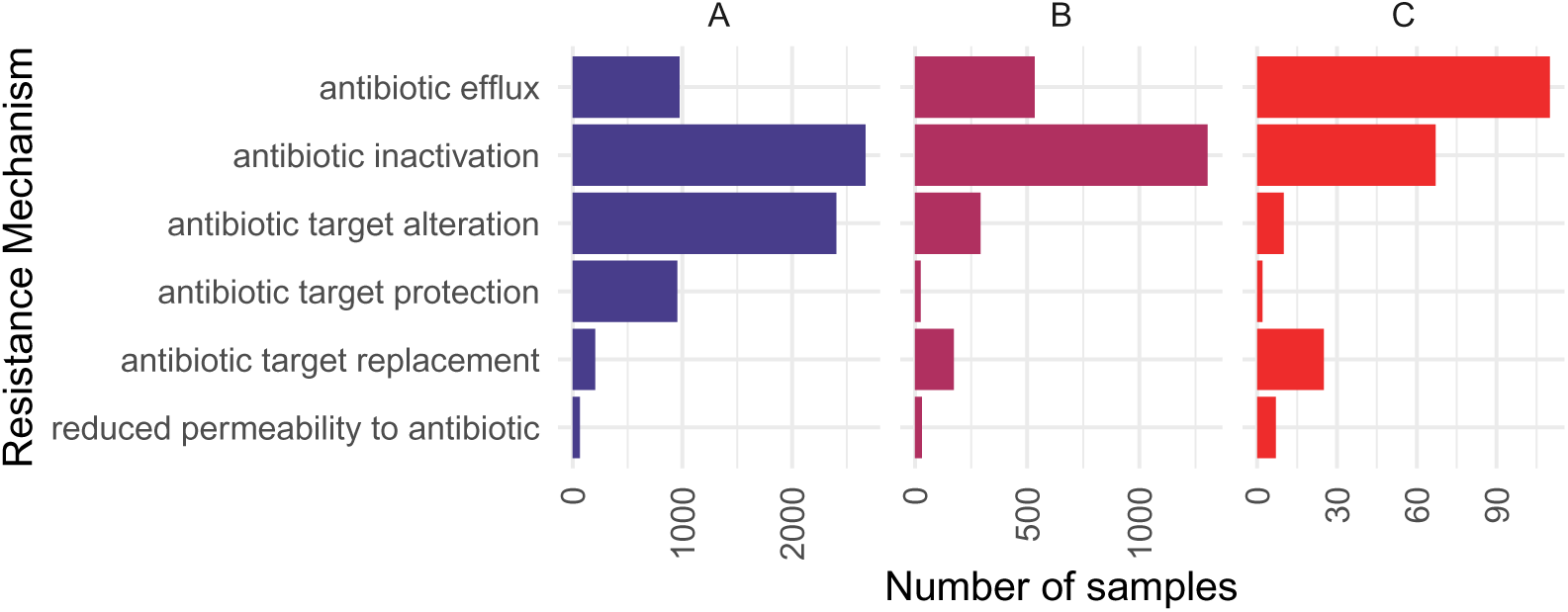
Antimicrobial resistance mechanism abundances of ARGs detected in all metagenomic samples (A), antimicrobial resistance mechanism abundances of higher public health risk ARGs detected in any metagenomic samples (B) and antimicrobial resistance mechanism abundances of higher public health risk ARGs in ESKAPE pathogens detected in the metagenomic samples (C). The number of occasions when ARGs with the given antimicrobial resistance mechanisms were detected, is presented on the horizontal axis.

### Basic characteristics, physical traits and behavioural characteristics associated with ARGs

The proportions of statistically significant and nonsignificant results of ARG abundances and traits or answers of the surveys have differed in the examined animal groups. The proportions of questions with significant (*p ≤* 0.05) and nonsignificant statistical test results by question types and by approaches (’A’, ’B’ or ’C’) are presented on Figure 5.

**Figure 5.**
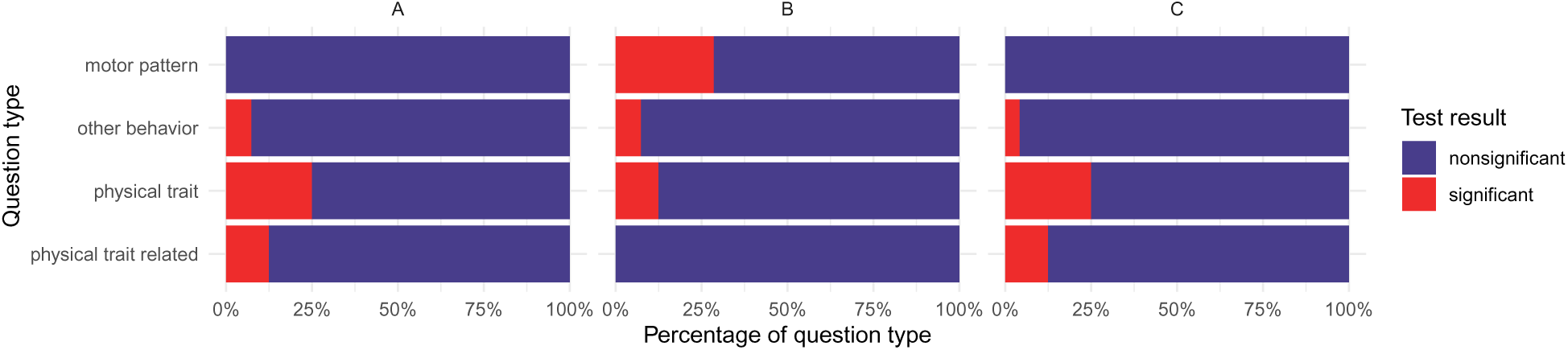
Proportions of significant (p ≤ 0.05) and nonsignificant associations by question groups and approaches (ARGs detected in all metagenomic samples (A), higher public health risk ARGs detected in any metagenomic samples (B) ESKAPE pathogen-related higher public health risk ARGs in the metagenomic samples (C)).

### Basic characteristics and physical traits

Analysing the associations of the numbers of ARG-positive and ARG negative samples in the light of basic characteristics, results represented in Table 1 were obtained. Considering all approaches, three out of six questions had statistically significant results.

**Table 1.** The number of samples containing ARGs (ARG+) and lacking ARGs (ARG-) associated with the following basic characteristics traits of dogs: head shape, origin, sex, sterilization status, breed, environment. ’Group 1’ and ’Group 2’ represent the order of traits indicated in the ’Characteristic’ column of the table. In column ’Approach’, ’A’ indicates ARGs detected in any canine metagenomic samples, ’B’ is for higher public health risk ARGs detected in any canine metagenomic samples and ’C’ stands for higher public health risk ARGs detected in ESKAPE pathogens.

| Characteristic<br>Group 1 / Group 2 | Approach | Number of samples |  |  |  | OR (95%CI) | p-value |
| --- | --- | --- | --- | --- | --- | --- | --- |
|  |  | Group 1 |  | Group 2 |  |  |  |
|  |  | ARG+ | ARG- | ARG+ | ARG- |  |  |
| Brachycephal / doliocephal, mesocephal | A | 71 | 12 | 606 | 50 | 0.49 (0.24 - 1.06) | 0.055* |
|  | B | 44 | 39 | 370 | 286 | 0.87 (0.54 - 1.42) | 0.560 |
|  | C | 7 | 105 | 29 | 598 | 1.37 (0.50 - 3.31) | 0.473 |
| Breeder / rescue, shelter | A | 286 | 28 | 1013 | 86 | 0.87 (0.55 - 1.41) | 0.557 |
|  | B | 172 | 142 | 628 | 472 | 0.91 (0.70 - 1.18) | 0.478 |
|  | C | 13 | 301 | 33 | 1067 | 1.40 (0.67 - 2.77) | 0.366 |
| Females / males | A | 827 | 68 | 852 | 83 | 1.18 (0.84 - 1.68) | 0.350 |
|  | B | 511 | 384 | 519 | 416 | 1.13 (0.76 - 1.70) | 0.561 |
|  | C | 27 | 828 | 33 | 861 | 0.85 (0.49 - 1.47) | 0.600 |
| Intact / sterilized | A | 87 | 10 | 1515 | 131 | 0.75 (0.38 - 1.66) | 0.440 |
|  | B | 44 | 53 | 938 | 709 | 0.63 (0.41 - 0.97) | 0.027* |
|  | C | 2 | 95 | 58 | 1589 | 0.58 (0.07 - 2.24) | 0.771 |
| Purebred / mutt | A | 677 | 62 | 935 | 80 | 0.93 (0.65 - 1.34) | 0.723 |
|  | B | 414 | 325 | 577 | 439 | 0.97 (0.80 - 1.18) | 0.770 |
|  | C | 36 | 703 | 25 | 991 | 2.03 (1.17 - 3.56) | 0.008** |
| Rural, suburban / urban | A | 575 | 46 | 837 | 76 | 1.13 (0.76 - 1.70) | 0.564 |
|  | B | 346 | 275 | 515 | 399 | 0.97 (0.79 - 1.20) | 0.834 |
|  | C | 23 | 598 | 31 | 883 | 1.10 (0.60 - 1.96) | 0.779 |

Associations of the ARG content of the saliva samples and physical traits, such as the size, fur color, fur type, tail type, ear shape, eye characteristics, fur length and fur texture are presented in Table 2. By certain questions, multiple comparisons of the trait options were performed.

**Table 2.** The number of samples containing ARGs (ARG+) and lacking ARGs (ARG-) associated with the questions regarding certain physical traits of dogs involved in the survey. ’Group 1’ and ’Group 2’ represent the order of traits indicated in the ’Trait’ column of the table. In column ’Approach’, ’A’ indicates ARGs detected in any canine metagenomic samples, ’B’ is for higher public health risk ARGs detected in any canine metagenomic samples and ’C’ stands for higher public health risk ARGs detected in ESKAPE pathogens.

| Trait | Approach | Number of samples |  |  |  | OR (95%CI) | p-value |
| --- | --- | --- | --- | --- | --- | --- | --- |
| Group 1 / Group 2 |  | Group 1 |  | Group 2 |  |  |  |
|  |  | ARG+ | ARG- | ARG+ | ARG- |  |  |
| **When the dog is standing next to someone of average height, how high are his shoulders?** |  |  |  |  |  |  |  |
| Under knee / knee high and above | A | 503 | 37 | 1035 | 95 | 1.25 (0.83 - 1.91) | 0.288 |
|  | B | 329 | 212 | 609 | 521 | 1.33 (1.07 - 1.65) | 0.008** |
|  | C | 15 | 526 | 45 | 1085 | 0.69 (0.35 - 1.27) | 0.261 |
| **What color is the dog?** |  |  |  |  |  |  |  |
| Whites / other colors | A | 795 | 54 | 685 | 70 | 1.50 (1.02 - 2.22) | 0.031* |
|  | B | 496 | 353 | 405 | 351 | 1.22 (0.99 - 1.49) | 0.056 |
|  | C | 36 | 813 | 21 | 735 | 1.55 (0.87 - 2.82) | 0.137 |
| Blacks / other colors | A | 149 | 11 | 1331 | 113 | 1.15 (0.60 - 2.42) | 0.757 |
|  | B | 90 | 71 | 811 | 633 | 0.99 (0.70 - 1.39) | 1.000 |
|  | C | 5 | 156 | 52 | 1392 | 0.86 (0.26 - 2.18) | 1.000 |
| Blacks / whites | A | 149 | 11 | 795 | 54 | 0.92 (0.46 - 2.00) | 0.860 |
|  | B | 90 | 71 | 496 | 353 | 0.90 (0.63 - 1.29) | 0.601 |
|  | C | 5 | 156 | 36 | 813 | 0.72 (0.22 - 1.89) | 0.664 |
| Whites / blacks, browns, reds, yellows | A | 795 | 54 | 543 | 58 | 1.57 (1.05 - 2.36) | 0.022* |
|  | B | 496 | 353 | 324 | 278 | 1.21 (0.97 - 1.50) | 0.086 |
|  | C | 36 | 813 | 18 | 584 | 1.44 (0.79 - 2.71) | 0.260 |
| Whites, merles / blacks, browns, reds, yellows | A | 816 | 55 | 543 | 58 | 1.58 (1.06 - 2.37) | 0.022* |
|  | B | 507 | 364 | 324 | 278 | 1.19 (0.96 - 1.48) | 0.098 |
|  | C | 36 | 835 | 18 | 584 | 1.40 (0.76 - 2.64) | 0.263 |
| Whites, merles, grays / blacks, browns, reds, yellows | A | 865 | 56 | 543 | 58 | 1.65 (1.10 - 2.47) | 0.012* |
|  | B | 531 | 390 | 324 | 278 | 1.17 (0.94 - 1.44) | 0.154 |
|  | C | 36 | 885 | 18 | 584 | 1.32 (0.72 - 2.49) | 0.396 |
| **How much white fur does dog have?** |  |  |  |  |  |  |  |
| All whites / others | A | 194 | 25 | 1341 | 107 | 0.62 (0.39 - 1.03) | 0.044* |
|  | B | 130 | 89 | 806 | 643 | 1.17 (0.86 - 1.57) | 0.307 |
|  | C | 7 | 212 | 53 | 1396 | 0.87 (0.33 - 1.95) | 0.847 |
| Almost all whites / others | A | 310 | 33 | 1225 | 99 | 0.76 (0.50 - 1.19) | 0.216 |
|  | B | 208 | 135 | 728 | 597 | 1.26 (0.99 - 1.62) | 0.059 |
|  | C | 13 | 330 | 47 | 1278 | 1.07 (0.53 - 2.04) | 0.871 |
| No white / any white | A | 670 | 55 | 865 | 77 | 1.08 (0.75 - 1.59) | 0.715 |
|  | B | 402 | 324 | 534 | 408 | 0.95 (0.78 - 1.16) | 0.619 |
|  | C | 21 | 705 | 39 | 903 | 0.69 (0.38 - 1.21) | 0.187 |
| All white / no white | A | 194 | 25 | 670 | 55 | 0.64 (0.38 - 1.10) | 0.095 |
|  | B | 130 | 89 | 402 | 324 | 1.18 (0.86 - 1.62) | 0.313 |
|  | C | 7 | 212 | 21 | 705 | 1.11 (0.39 - 2.75) | 0.821 |
| **Is dog's tail curly?** |  |  |  |  |  |  |  |
| Not curly / curly | A | 1036 | 93 | 396 | 32 | 0.90 (0.57 - 1.38) | 0.677 |
|  | B | 625 | 505 | 242 | 186 | 0.95 (0.76 - 1.20) | 0.689 |
|  | C | 37 | 1093 | 16 | 412 | 0.87 (0.47 - 1.70) | 0.641 |
| **What is the dog's ear shape like?** |  |  |  |  |  |  |  |
| Long pendant ears / others | A | 181 | 14 | 1355 | 118 | 1.13 (0.63 - 2.17) | 0.779 |
|  | B | 119 | 76 | 818 | 656 | 1.26 (0.92 - 1.73) | 0.146 |
|  | C | 9 | 186 | 50 | 1424 | 1.38 (0.59 - 2.89) | 0.406 |
| Surgically cropped ears / others | A | 18 | 2 | 1518 | 130 | 0.77 (0.18 - 6.92) | 0.669 |
|  | B | 10 | 10 | 927 | 722 | 0.78 (0.29 - 2.10) | 0.653 |
|  | C | 1 | 19 | 58 | 1591 | 1.44 (0.03 - 9.40) | 0.515 |
| **Are dog's eyes different colors?** |  |  |  |  |  |  |  |
| Not different eyes / different eyes | A | 1476 | 126 | 50 | 3 | 0.70 (0.14 - 2.23) | 0.794 |
|  | B | 900 | 703 | 32 | 21 | 0.84 (0.46 - 1.52) | 0.576 |
|  | C | 54 | 1549 | 6 | 47 | 0.27 (0.11 - 0.82) | 0.011* |
| **How long is dog's fur on his back and sides?** |  |  |  |  |  |  |  |
| Long / short | A | 216 | 13 | 781 | 74 | 1.57 (0.85 - 3.15) | 0.170 |
|  | B | 134 | 95 | 479 | 376 | 1.11 (0.82 - 1.51) | 0.548 |
|  | C | 15 | 214 | 26 | 829 | 2.23 (1.08 - 4.47) | 0.019* |
| Long / medium, short | A | 216 | 13 | 1296 | 118 | 1.51 (0.83 - 2.98) | 0.189 |
|  | B | 134 | 95 | 787 | 628 | 1.13 (0.84 - 1.51) | 0.431 |
|  | C | 15 | 214 | 43 | 1372 | 2.24 (1.13 - 4.19) | 0.012* |
| **Does the dog have soft fur or rough and bristly fur?** |  |  |  |  |  |  |  |
| Rough(wiry) / soft | A | 220 | 17 | 1205 | 109 | 1.17 (0.68 - 2.12) | 0.698 |
|  | B | 142 | 95 | 735 | 580 | 1.18 (0.88 - 1.58) | 0.256 |
|  | C | 6 | 231 | 51 | 1264 | 0.64 (0.22 - 1.52) | 0.451 |

### Behavioral traits

Due to the high number of behavioural traits included in the original questionnaire, only the most significant results are mentioned in the Results section. See Supplementary table 7-9. for full behavioral result tables.

By questions that could have been answered with Never or Not never (Always, Often, Sometimes, Rarely), two statistically significant results were found. Statistical test results for all questions of this answer category are presented in Table 3.

**Table 3.** The number of samples containing ARGs (ARG+) and lacking ARGs (ARG-) associated with questions regarding the behaviour of dogs involved in the survey. ’Group 1’ and ’Group 2’ represent answer ’Never’ and ’Not never’ (Always, Often, Sometimes, Rarely), respectively. In column ’Approach’, ’A’ indicates ARGs detected in any canine metagenomic samples, ’B’ is for higher public health risk ARGs detected in any canine metagenomic samples and ’C’ stands for higher public health risk ARGs detected in ESKAPE pathogens.

| Question | Approach | Number of samples |  |  |  | OR (95%CI) | p-value |
| --- | --- | --- | --- | --- | --- | --- | --- |
|  |  | Never |  | Not never |  |  |  |
|  |  | ARG+ | ARG- | ARG+ | ARG- |  |  |
| Dog walks away or avoids being patted | A | 705 | 50 | 841 | 80 | 1.34 (0.92 - 1.98) | 0.120 |
|  | B | 422 | 333 | 521 | 401 | 0.98 (0.80 - 1.19) | 0.805 |
|  | C | 36 | 719 | 24 | 898 | 1.87 (1.08 - 3.31) | 0.024* |
| Dog shows barrier aggression | A | 1037 | 82 | 507 | 50 | 1.25 (0.85 - 1.83) | 0.249 |
|  | B | 634 | 486 | 307 | 250 | 1.06 (0.86 - 1.31) | 0.566 |
|  | C | 43 | 1077 | 17 | 540 | 1.27 (0.70 - 2.39) | 0.486 |
| Dog lays in one place all day long | A | 734 | 68 | 815 | 64 | 0.85 (0.58 - 1.23) | 0.366 |
|  | B | 434 | 368 | 513 | 367 | 0.84 (0.69 - 1.03) | 0.085 |
|  | C | 28 | 774 | 31 | 849 | 0.99 (0.57 - 1.72) | 1.000 |
| Dog tilts his head | A | 140 | 20 | 1384 | 110 | 0.56 (0.33 - 0.98) | 0.029* |
|  | B | 92 | 68 | 838 | 656 | 1.06 (0.75 - 1.50) | 0.801 |
|  | C | 5 | 155 | 53 | 1441 | 0.88 (0.27 - 2.23) | 1.000 |

By the comparison of the number of ARG-positive and negative samples by questions with answer groups Agree (Strongly agree, Agree) / Disagree (Strongly disagree, disagree) several statistically significant results were obtained. Statistical test results for all questions, including those with an unclear biomedical significance are presented in Table 4.

**Table 4.**
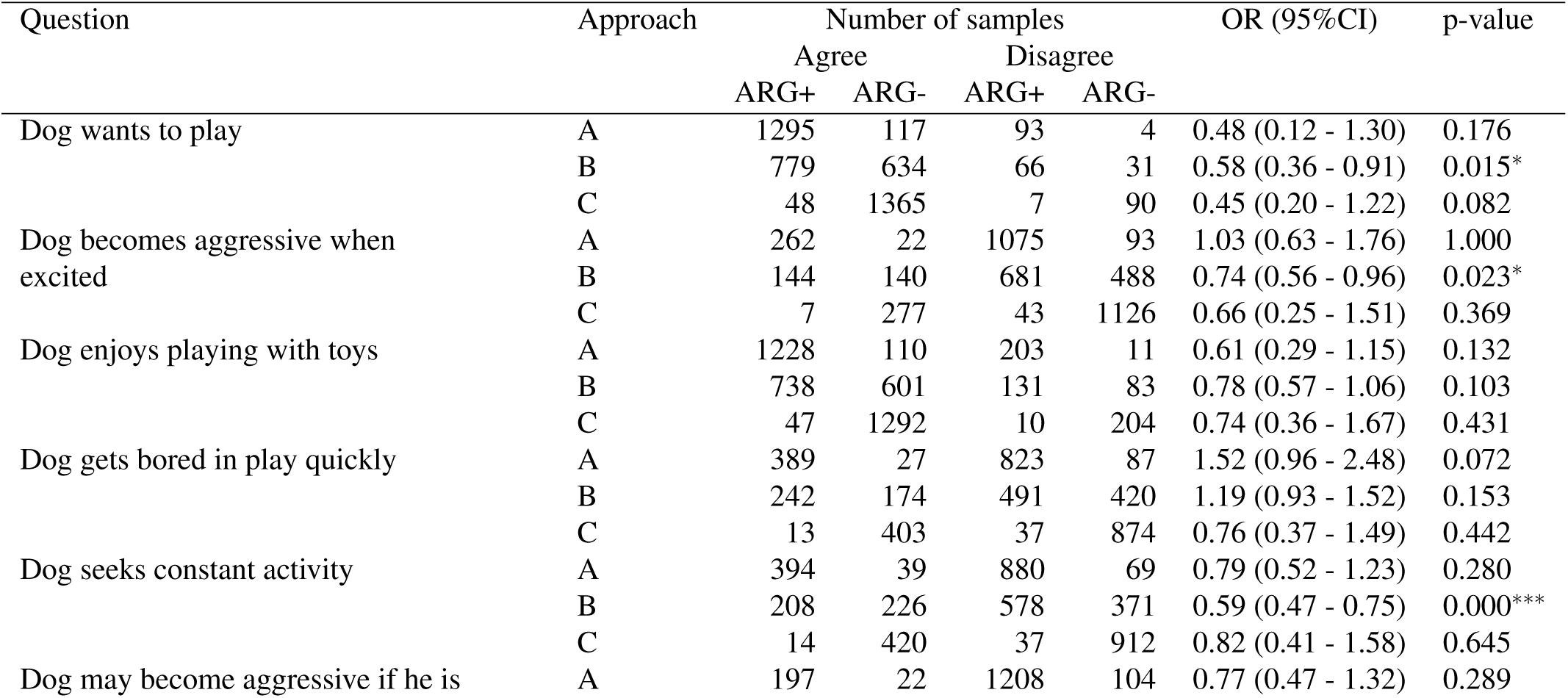
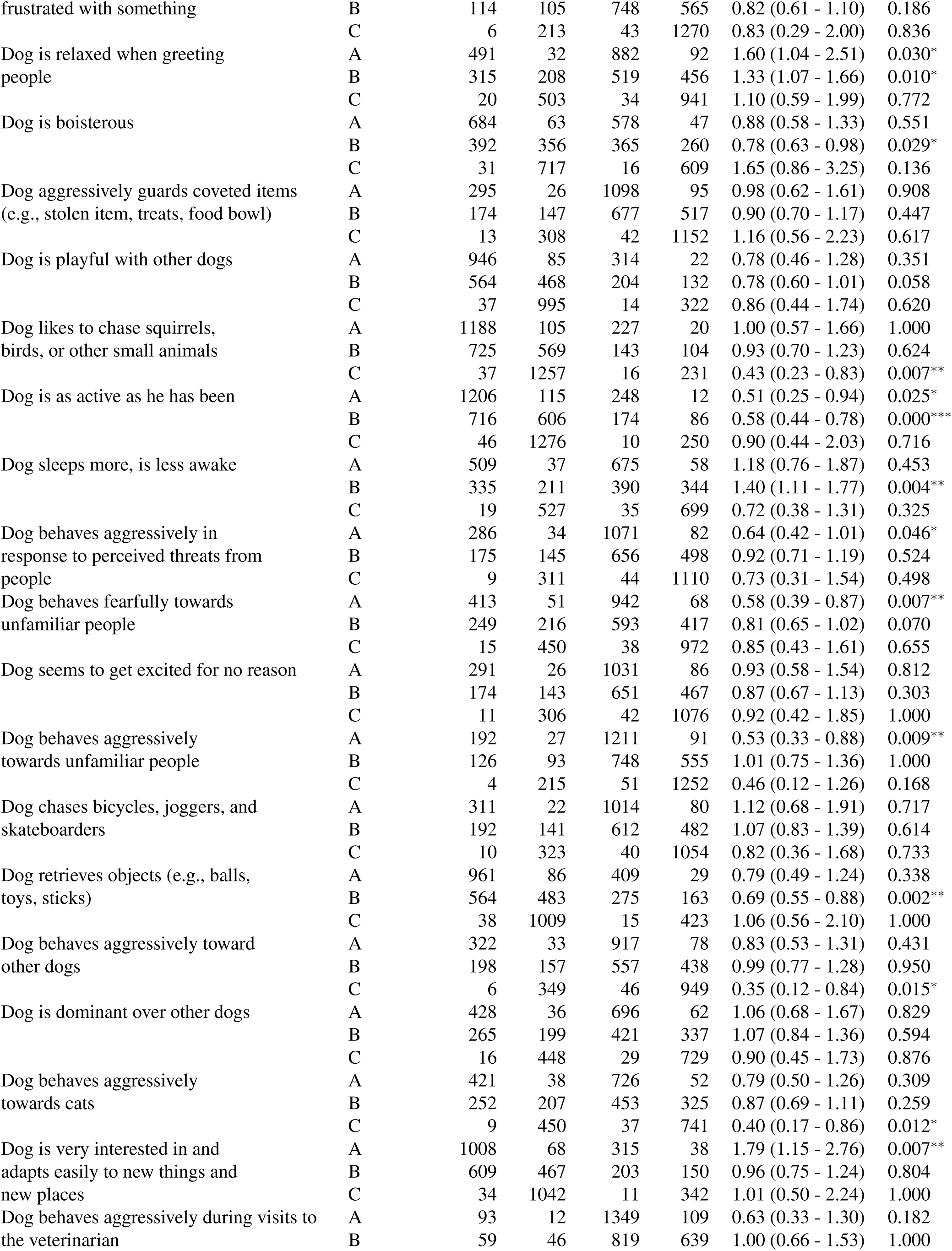
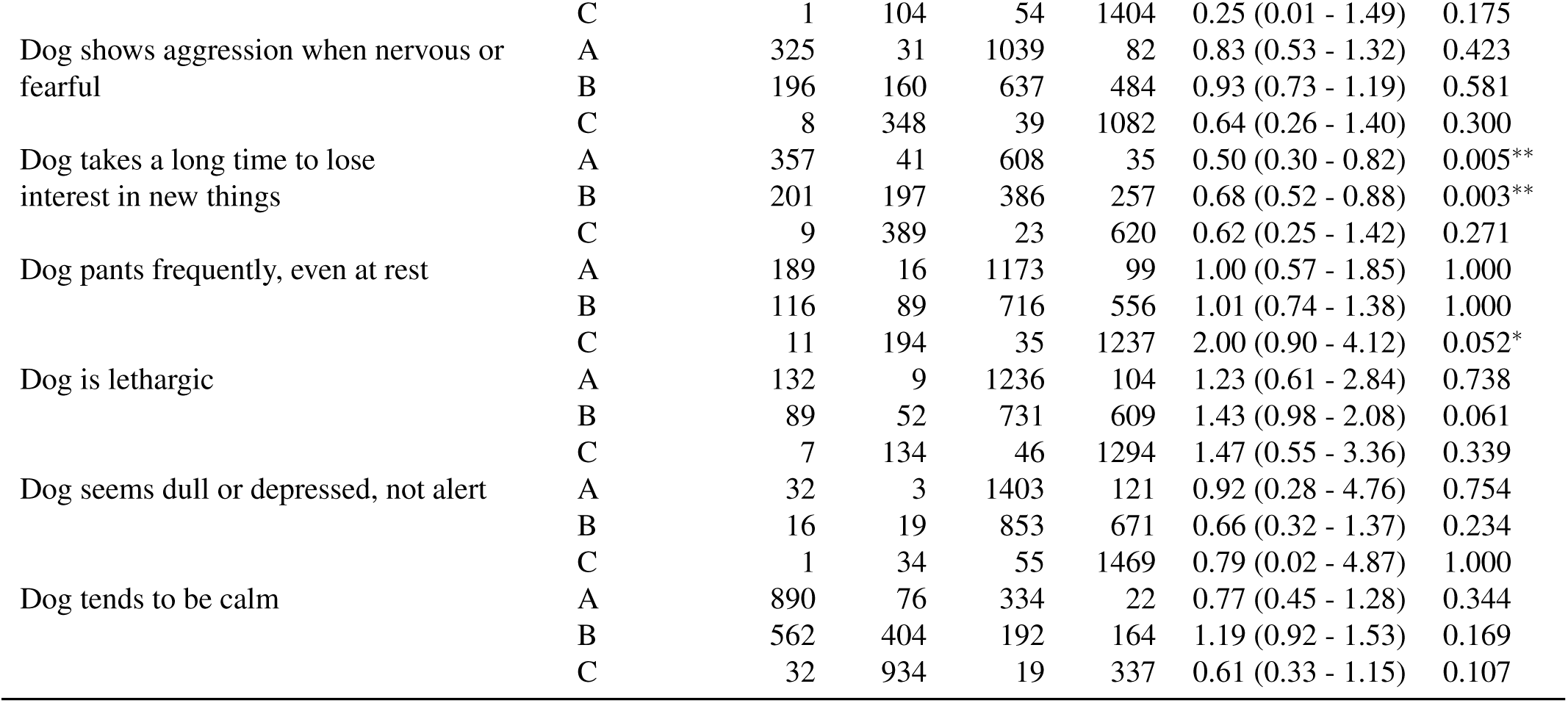
The number of samples containing ARGs (ARG+) and lacking ARGs (ARG-) associated with questions regarding the behaviour of dogs involved in the survey. ’Group 1’ and ’Group 2’ represent answer ’Agree’ (Strongly agree, Agree) and ’Disagree’ (Strongly disagree, disagree), respectively. In column ’Approach’, ’A’ indicates ARGs detected in any canine metagenomic samples, ’B’ is for higher public health risk ARGs detected in any canine metagenomic samples and ’C’ stands for higher public health risk ARGs detected in ESKAPE pathogens.

Comparing answer groups of Agree (Strongly agree, Agree) / Neither agree, nor disagree / Disagree (Strongly disagree, disagree) gave a similar but not exactly identical results that can be found in Supplementary Table 9.

## Discussion

Throughout the study, several AMR-related aspects of the canine oral cavity and saliva have been revealed.

All ARG-associated bacterial genera identified in the canine salivary samples have been previously described in the canine oral cavity (*Porphyromonas spp., Pasteurella spp., Frederiksenia spp., Pseudomonas spp., Conchiformibius spp., Bacteroides spp., Glaesserella spp., Riemerella spp., Escherichia spp., Prevotella spp., Capnocytophaga spp., Klebsiella spp., Mannheimia spp., Histophilus spp., Streptococcus spp., Serratia spp., Haemophilus spp., Enterobacter spp., Clostridioides spp., Parabacteroides spp.*)^15–23^, or along the canine gastrointestinal tract (*Alistipes spp., Lelliottia spp.*).^24,25^ Wielerella, a genus of Neisseriaceae family has not been previously associated with the canine oral cavity according to our best knowledge.

After the bioinformatic analysis conducted based on reference data from the Comprehensive Antimicrobial Resistance Database (CARD) a high number of AMR determinants hits were identified in the canine saliva samples. AMR determinants stored in CARD are all associated with AMR.^26^ Several genes are directly responsible for the appearance of AMR (e.g. *dfra14, dfra20, sul1, sul2, SRT-2, SRT-3, SAT-4, tet* family, *ROB* family, *APH* family, OXA family etc.).^26^ However, several ARGs mentioned in the results section are essential for cell viability, and eventually also cause AMR. Certain ARGs (e.g. *mel, msbA*) are sequence homologs of previously known ARGs that can be responsible for AMR themselves^27,28^, and some encode for intrinsic, non-transferable AMR characteristics (e.g. *mexA, mexB, mexC, mexE*.^29^ In other cases, the mutation of an intrinsic, originally non-AMR encoding gene causes specific antibiotic effects (e.g. *Escherichia coli EF-Tu* mutants conferring resistance to Pulvomycin, *Escherichia coli UhpT* with mutation conferring resistance to fosfomycin). Other genes that indirectly induce or regulate AMR facilitators (e.g. *ArmR, baeR, H-NS, marA, ramA*)^30–34^ are also inevitable in the phenotypical appearance of AMR. Furthermore, genes encoding the members of complexes that realize AMR or broader spectrum AMR were also detected (e.g. *mdtB, mdtC, mex* family).^35–37^ In case of the above mentioned mdtBC multidrug transporter, even the regulator, *baeR* was identified in many samples.^31^ At other cases only subunits or functional cooperators of AMR determinants complexes (e.g. efflux pumps) were identified (e.g. *qacL, oprM, oprN, OpmB, opmE*)^38–41^. Even though being associated to AMR, the presence of certain determinants, such as *CRP, nalC, nalD* is even beneficial due to the repression or modulation of multidrug resistance. Despite the presence of *CRP, nalC* and *nalD* both in the CARD database and in numerous samples, these genes were left out of further analysis steps due to their anti-AMR effect.^42–44^ Even though, AMR determinants are present in the canine salivary samples, their occurrence does not necessarily lead to the phenotypical expression of AMR. Furthermore, from a One Health point of view, those ARGs have a special significance which can spread via horizontal gene transfer among bacteria. Nonetheless, ARGs which are intrinsic and are not escorted by mobile genetic elements can only be transferred within the (potentially pathogenic) bacteria that they were found at. To enhance the public health risk, Zhang and colleagues^13^ used four indicators (higher human accessibility, mobility, pathogenicity, and clinical availability) to identify higher public health risk ARGs. Based on the indicators, higher public health risk ARGs and higher public health risk ARGs detected in ESKAPE pathogens have a strong AMR potential and an enhanced public health significance.

However, another means of the evaluation of the public health importance of the detected ARGs is the assessment of the affected drug classes. A possible baseline for this is the WHO list of Critically important antimicrobials for human medicine^45^. According to this WHO review, Critically Important Antimicrobials (CIAs) (devided into Highest Priority Critically Important Antimicrobials (Category 1) and High Priority Critically Important Antimicrobials (Category 2)) are in need of the most urgent AMR risk management steps. Among the overall ARG set, the most hits may affect tetracyclines, cephalosporins, peptides and penams. Antibiotic compounds from these groups, such as amoxicillin–clavulanate, first-generation cephalosporins or doxycycline are the flagships of routine antibiotic administration in small animal veterinary practices.^46–48^ However, other compounds among that are Highest Priority CIAs, namely 3rd, 4th and 5th generation cephalosporins, glycopeptides (e.g. vancomycin) and polymyxins (polymyxin-B, colistin) are also the members of these groups. Members of the *van* gene family, that confer glycopeptide resistance^49^ and polimixin resistance associated genes including *eptB,* plasmid encoded *MCR-9.1^50^, mtrA, PmrF, AcrF, AcrE, KpnH, KpnF, KpnG, KpnE,*^51,52^ have also been identified in the canine saliva samples. However, 3rd and 4th generation cephalosporin resistance associated *CTX-M* gene family members were not found in the samples, determinants from other extended-spectrum beta-lactamase gene families, such as *SHV (SHV-1, SHV-11)* and *TEM (TEM-135, TEM-178)* were detected.^53^ A regulatory member of the Methicillin-Resistant *Staphylococcus aureus* and *Staphylococcus epidermidis* associated *mec* family (*mecI*) has also been identified.^54^ Regarding the remaining Highest Priority CIAs (Macrolides and ketolides, Quinolones), the number of resistance determinants against them was so high, that they were presented to become the second most and the most frequently affected drug classes considering higher public health risk ARGs and ESKAPE pathogen higher public health risk ARGs, respectively. High Priority CIAs aminoglycosides were the third most affected group by higher public health risk ARGs. Furthermore, 59 AMR determinant types appeared against carbapenems and other penems, that are also Category 2 CIAs.

The shift of relative abundances towards antibiotic efflux, a mechanism type capable of affecting multiple drug classes parallelly in ESKAPE pathogen-related higher health risk ARGs is because multidrug resistance exhibited by ESKAPE pathogens is widespread.^55^

Relatively more physical trait-related questions had significant results which may be associated with the fact that physical traits are objectively observable, while behavioural traits can be determined rather subjectively. While sex, early life (origin) and - despite the supposed lifestyle differences - environment do not determine the salivary ARG frequencies, other basic characteristics do. Interestingly, doliocephalic dogs are slightly more often ARG-positive than brachycephalics. In contrast, strong evidence of brachycephalic breeds being less healthy has been provided^56^ that could possibly lead to more visits to the veterinarian. However, popular brachycephalic breeds, such as the French bulldog have a shorter life expectancy^57^ that can contribute to the balance. According to a common belief, ’mutts are overall healthier than purebreds’. Considering certain inherited disorders, recently bred, similar canine lineages appear to be more susceptible.^58^ The higher number of ESKAPE pathogen-related higher health risk ARG-positive samples may either due to this or owner habits regarding the frequencies of veterinary visits having more expensive dogs. Nonetheless, Singleton and colleagues found that that antibiotic use was less likely in sterilized dogs^59^, higher health risk ARGs were significantly less frequently detected in intact canines. The reason might be that the owners of sterilized animals are more likely to seek veterinary care due to closer bonds or more responsibility towards the animals. The significantly higher number of higher public health risk ARGs under knee-high can be underlain by the phenomenon that toy/pet dogs have a closer bond to their owners and share the same environment. Regarding the colors, there seem to be a clear trend of more ARG-positive samples among white dogs than other colors, even though the question regarding the white fur ratio of the dogs suggests the contrary. The most possible reason for this inconsistency can be seen on Supplementary figure 6 and 7 where the breed-color and the breed-whiteness associations are presented. The majority positive of answers to the question regarding the white fur ratio of the dogs, and namely if the dog was all white arrived from the owners of golden retrievers and labrador retrievers (Supplementary figure 7). Even though, these dogs are often light, yellow coat color is far more frequent than pure white. By the questions regarding the exact color of the same breeds (Supplementary figure 6), few answered that their labradors or golden retrievers are white. Based on this, we can assume that the question regarding the coat whiteness ratio of dogs was considered to apply to the ratio of lightness fur to dark fur. However, there might be a difference in light fur colored dogs and pure white dogs. While labrador retrievers and golden retrievers are as or even more prone to allergies that might lead to skin infections as several breeds appearing among the white fur color dogs^60^, the genetic background of the white coat is different from that of light yellow.^61^ Genes responsible for white fur also cause severe phenotypes, including extensive depigmentation, hearing loss, and acute eye and bone disorders^61^ that may facilitate more frequent veterinary visits. The finding, that a shift to ESKAPE pathogen-related high risk ARG-positive samples also appeared in dogs with heterochromia is in line with the findings of white fur color. Breeds with the highest number of answers of white fur color (american pit bull terriers, border collies, huskies etc.) are also prone to having different eye colors.**^?^**oreover, also when white, merle or gray color dogs were examined together, higher ARG-positivity was detected. In merle and grey dogs, heterochromia is also common^62^ and certain hereditary diseases are also more prevalent (ocular and auditory anomalies).^63,64^ Even though, long pendant ears may facilitate external ear canal infections, no related significant effect could have been observed in the number of ARG-positive samples. In contrast, a rise in the number of ESKAPE pathogen-related higher health risk ARG-positives among canines with long fur was noticeable.

The evaluation of behavioral findings is very difficult. Dogs that walk away or avoid being patted were statistically significantly more prone to have higher public health ARGs related to ESKAPE pathogens, while dogs that tilt their head carried less ARGs in general. Significantly more ARG-positive samples appeared among canines that are relaxed when greeting people or are very interested in and adapt easily to new things and new places than among those that are not. At the same time, more ARG-positive saliva samples were found in dogs that are not as active as they have been, do not behave fearfully or aggressively towards unfamiliar people, do not behave aggressively in response to perceived threats from people or do not take a long time to lose interest in new things than in those that do. Dogs that are relaxed when greeting people or sleep more and be less awake contained higher public health risk ARGs more frequently than those that are not. In contrast, dogs that do not want to play, do not become aggressive when excited, do not seek constant activity, are not boisterous, do not retrieve objects (e.g. balls, toys, sticks), are not as active as they have been or do not take a long time to lose interest in new things were more often higher public health risk ARG-positive than those that do. The number of samples containing higher health risk ARGs in ESKAPE pathogens was higher in dogs that pant frequently even at rest than in those than do not, but canines that do not like to chase squirrels, birds or other small animals or do not behave aggressively toward other dogs or cats than those that do. Dogs that are relaxed when greeting people, that are not as active as they have been or that do not take a long time to lose interest in new things are both more often ARG-positive and public health risk ARG-positive than dogs behaving in the opposite ways. A pattern that seems to appear considering all significant results is that dogs that are more active less often harbored ARGs, higher health risk ARGs or ESKAPE pathogen-related higher health risk ARGs in their saliva samples. All in all, this observation applied to the followings. Dogs that are relaxed in general, are relaxed when greeting people, sleep more and be less awake, are not boisterous, are not as active as they have been, do not want to play, do not seek constant activity, do not retrieve objects (e.g. balls, toys, sticks), do not like to chase squirrels, birds or other small animals, do not take a long time to lose interest in new things or are very interested in and adapt easily to new thing and new places were in higher numbers associated with ARG, higher health risk ARG or ESKAPE pathogen higher health risk ARG-positivity. However, other activity-related questions (e.g., dog lays in one place all day long, dog seems to get excited for no reason, dog chases bicycles, joggers, and skateboarders, dog is lethargic, dog seems dull or depressed, not alert, dog tends to be calm) did seem to be in line with this tendency and had no statistically significant results by any approaches. Furthermore, another possible trend is that dogs that are less aggressive in general, namely by not behaving fearfully or aggressively towards unfamiliar people, not behaving aggressively in response to perceived threats from people, not becoming aggressive when excited are more often ARG or higher health risk ARG-positive. The explanation this phenonemon is unclear, but might derive from the fact that less aggressive dogs are more favorable as pets, and due to the closer human bonds are brought to veterinarians more regularly. On the other hand, some questions that reported about the aggression of canines (e.g. dog shows barrier aggression, dog may become aggressive if he is frustrated with something, dog aggressively guards coveted items (e.g., stolen item, treats, food bowl), dog is dominant over other dogs, dog behaves aggressively during visits to the veterinarian, dog shows aggression when nervous or fearful) had no statistically significant results.

Interestingly, the proportion of statistically significant results was the highest by ’motor pattern’ of all question types by approach ’B’, while no significant results of this question type appeared by any other approaches (’A’ or ’C’). Questions with statistically significant results of this type aimed to reveal the playfulness of dogs (dog wants to play, dog retrieves objects (e.g. balls, toys, sticks)). According to this, decreased playfulness might be especially characteristic for elevated higher public health risk ARG presence in canine salivary bacteria. In contrast to this, no significant elevation in the presence of higher public health risk ARGs was observable by the following playfulness-related questions: dog enjoys playing with toys, dog gets bored in play quickly, dog is playful with other dogs. By approach ’A’ and ’C’, question type ’physical trait’ had the highest proportion of significant test results. This finding may be associated to the fact, the physical traits are easier to objectively observe than behavioural traits that can be evaluated by the owners rather subjectively.

To fully understand the One Health implications of the results arisen, further studies assessing the relatedness of the skin microbiome of pets and their owners would be required. However, disturbing findings such as the high detection rate of peptides and other CIAs indeed draw attention to the potential One Health and public health significance of human-pet proximity.

## Methods

### Data

Canine salivary genomic datasets were obtained from the National Center for Biotechnology Information (NCBI) Sequence Read Archive (SRA) repository. The BioProject used is stored under the Accession Number PRJNA675863 submitted by the Broad Institute, Darwin’s Ark project. The project contained the shotgun metagenomic sequencing data of canine saliva samples to investigate polymorphisms associated to complex canine traits.^14^ The BioProject contained short read data from 2215 canine saliva samples, out of which 1830 was used in the study. The selection process was based on the completeness of the metadata information of the samples. Metadata used derived from the Broad Institute’s Darwin’s Ark project survey dataset (https://datadryad.org/stash/dataset/doi:10.5061/dryad.g4f4qrfr0, accessed on 8/12/2023). Throughout the analysis steps, the study of the presence of any ARGs, higher public health risk ARGs and ESKAPE pathogen-related higher public health risk ARGs within the metagenomic samples was signed with ’A’, ’B’ and ’C’, respectively. The survey that gave the basis for the trait associated AMR studies was filled voluntarily by dog owners. Out of 12 questionnaires, 11 surveyed canine behaviours, and 1 focused on physical traits. Breed or suspected breeds, age, sex and sterilization status were collected initially.^14^ Samples were analysed based on their unique darwinsark_ids. However, in case of sex, only those BioSample identifiers were included in the analyses that could have been associated with unique darwinsark_id identifiers or with no darwinsark_id identifiers. All questions in the survey of Morrill and colleagues^14^ have been included in the analysis workflow. However, only questions with possible biomedical explanations are presented in the Results section. All other question results are among the Supplementary materials.

### Bioinformatic and statistical analysis

After quality control, the trimming and filtering of the raw short reads was performed with TrimGalore (v.0.6.6, https://github.com/FelixKrueger/TrimGalore, accessed on 8/12/2023), with a quality treshold of 20. More than 50 bp long reads were taxonomically classified using Kraken2 (v2.1.1)^65^ using a database created (4/6/2022) based on the NCBI RefSeq complete archaeal, bacterial, viral and plant genomes. For precise species assignment, a confidence parameter of 0.5 was used. The taxon classification results were processed in the R-environment^66^ using the packages phyloseq (v1.36.0)^67^ and microbiome (v1.14.0)^68^. Bacterial reads were assembled to contigs with MEGAHIT (v1.2.9)^69^ using default settings. Kraken2 was used again for the taxonomic classification of the contigs based on the same database as mentioned above. Possible open reading frames (ORFs) were extracted from the contigs with Prodigal (v2.6.3).^70^ Amino-acid sequences translated from the ORFs were ORFs were aligned to the reference ARG sequences of the Comprehensive Antibiotic Resistance Database (CARD, v.3.1.3)^71,72^ with Resistance Gene Identifier (RGI, v5.2.0) with Diamond.^73^ ORFs of ’Perfect’ or ’Strict’ classification were further filtered with 90% identity and 90% coverage to the reference sequences. The public health risk categorization of the ARGs was based on a publication of Zhang and colleagues.^13^ Zhang and colleagues^13^ used four indicators to categorize ARGs having higher public health risk: higher human accessibility, mobility, pathogenicity, and clinical availability. Human accessibility was based on the transfer likelihood of the ARG from the environment to the human microbiota. Mobility was defined by the relatedness of mobile genetic elements to the ARG. Human pathogenicity was based on appearance rates in human pathogens compared to that in non-pathogenic bacteria. Clinical availability was evaluated according to the use and clinical relevance of the affected antimicrobial agents. The independence of ARG frequencies on various grouping variables was tested using Fisher’s exact test.^74^ A p-value of < 0.05 was considered statistically significant. All further data management procedures, analyses, and plotting were performed in the R-environment (v4.2.1)^66^.

## Declarations

### Ethics approval and consent to participate

Not applicable.

### Consent for publication

Not applicable.

### Data Availability

The short-read data of samples are publicly available and accessible through the PRJNA675863 from the NCBI Sequence Read Archive (SRA).

## Author contributions statement

NS takes responsibility for the integrity of the data and the accuracy of the data analysis. AGT and NS conceived the concept of the study. AGT and NS participated in the bioinformatic analysis. AGT, IT, and NS participated in the drafting of the manuscript. AD, AGT, ÁVP, DLT, IT, LM, LR, NS, and TN carried out the critical revision of the manuscript for important intellectual content. All authors read and approved the final manuscript.

## Acknowledgements

The authors would like to thank the providers of BioProject PRJNA675863^14^ and, furthermore, Kathleen Morrill; Elinor Karlsson; Kerstin Lindblad-Toh, (Karlsson Laboratory, Broad Institute) and the Darwin’s Ark project (darwinsark.org, accessed on 17 January 2024) for providing the datasets and metadata.

## Authors’ information

Not provided.

## Supplementary materials

**Table 5.**
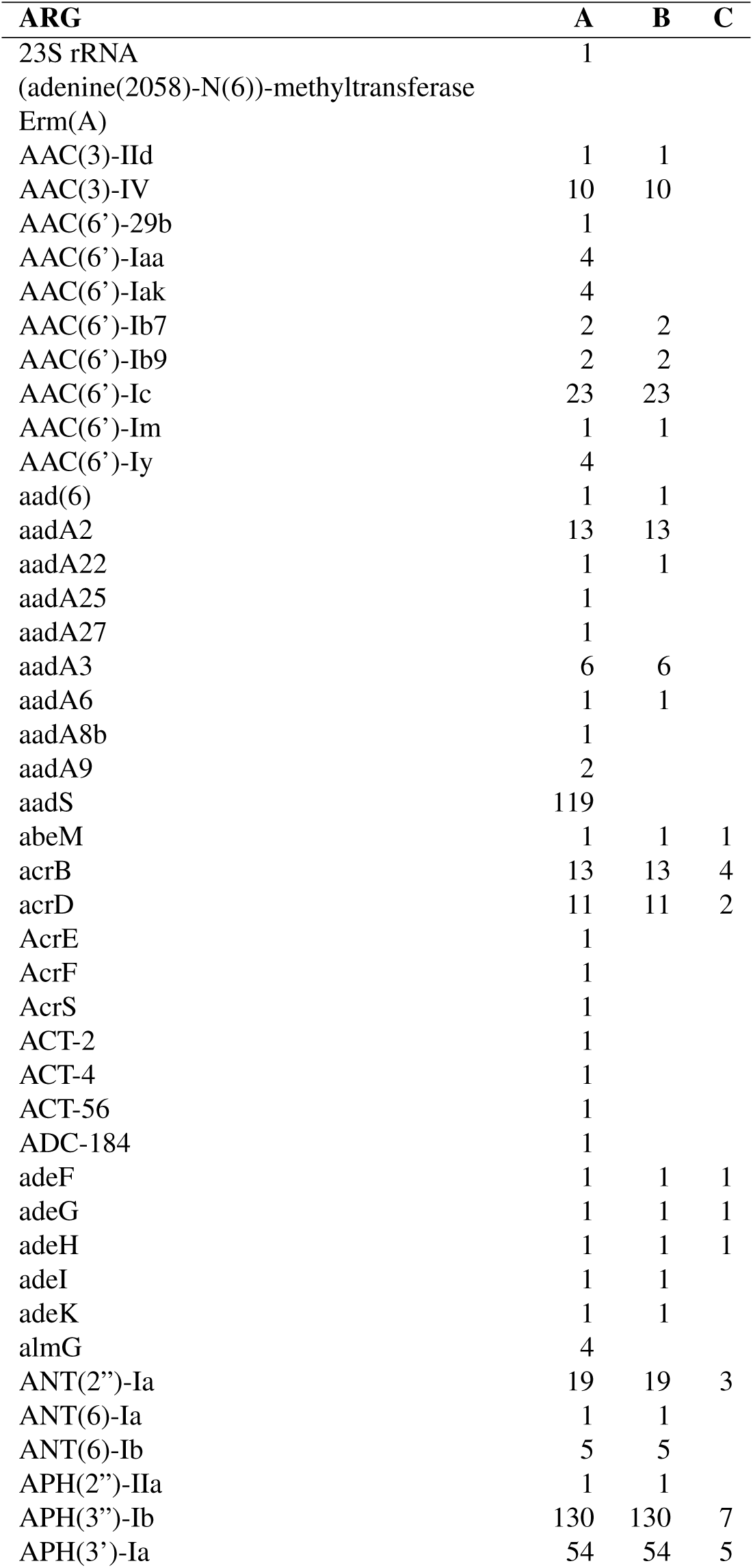
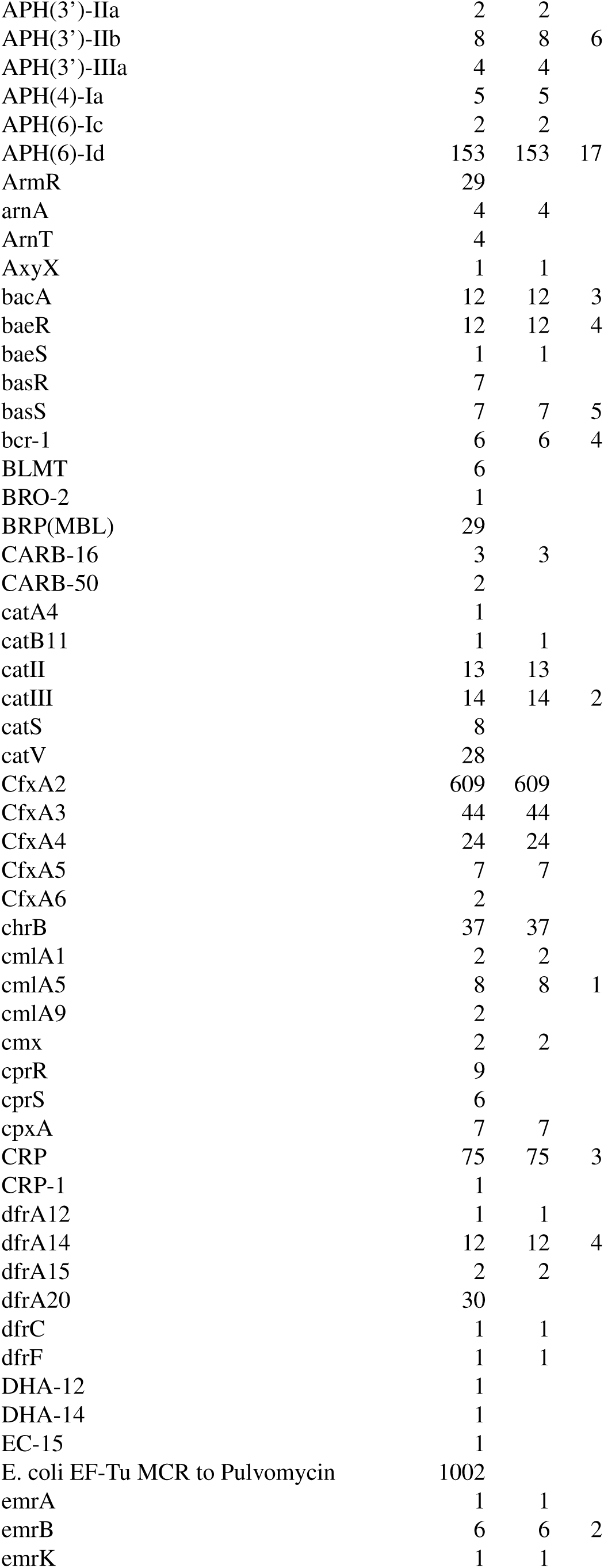
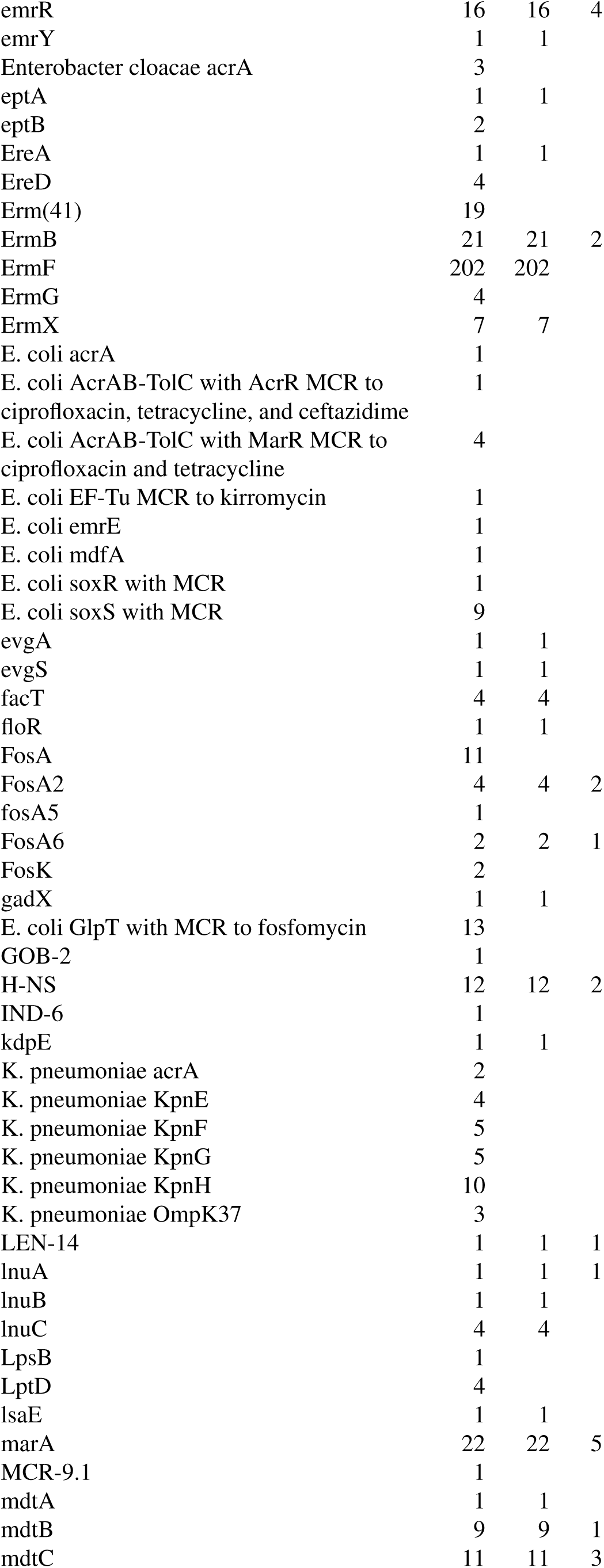
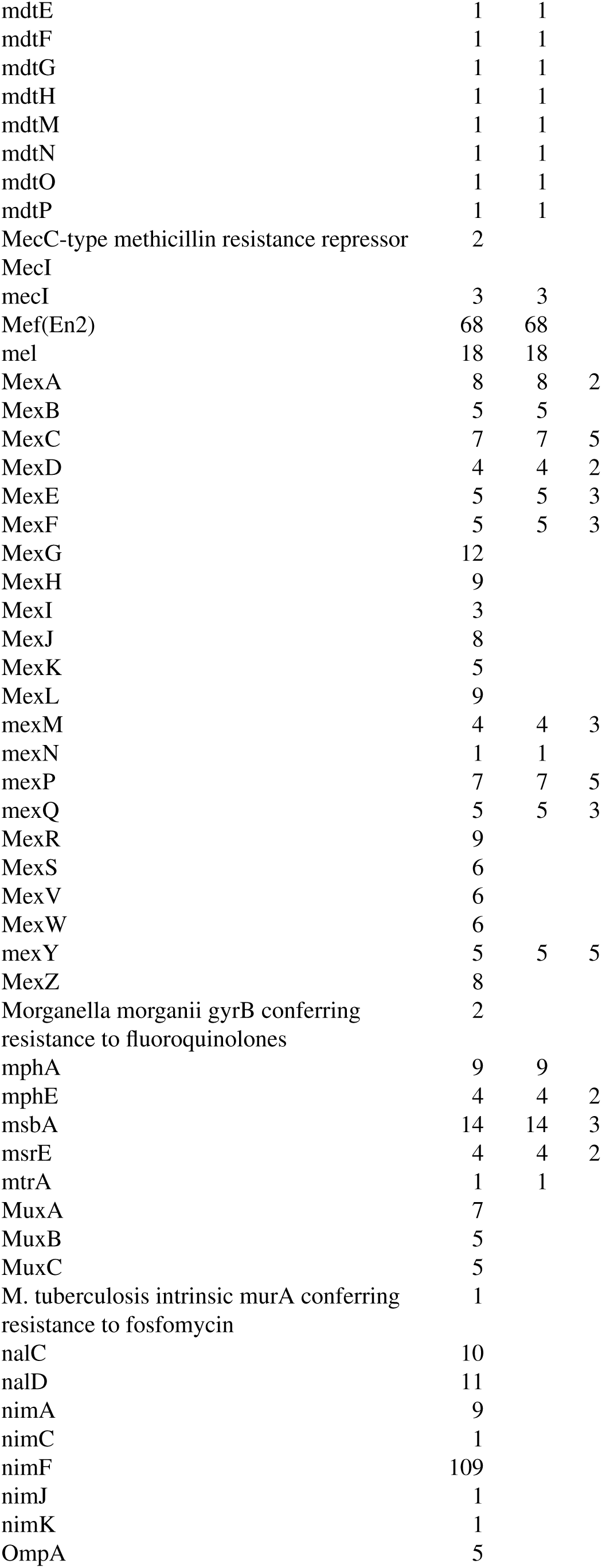
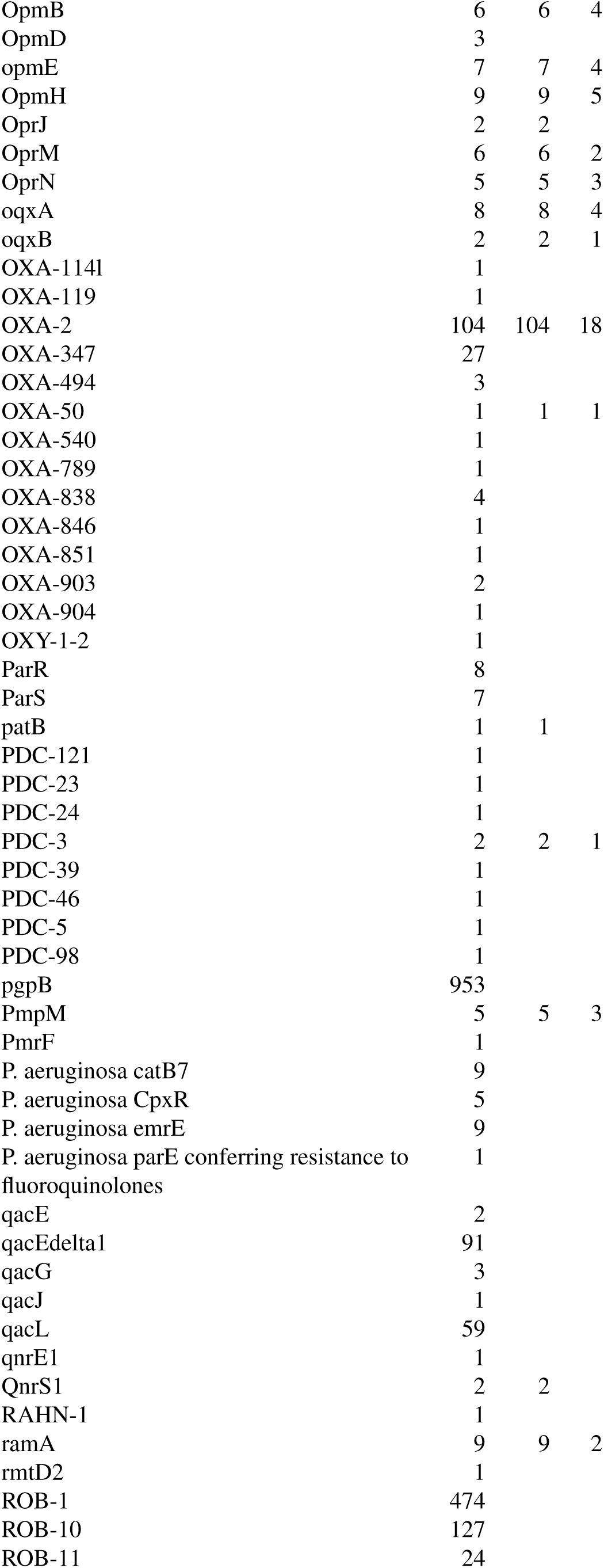
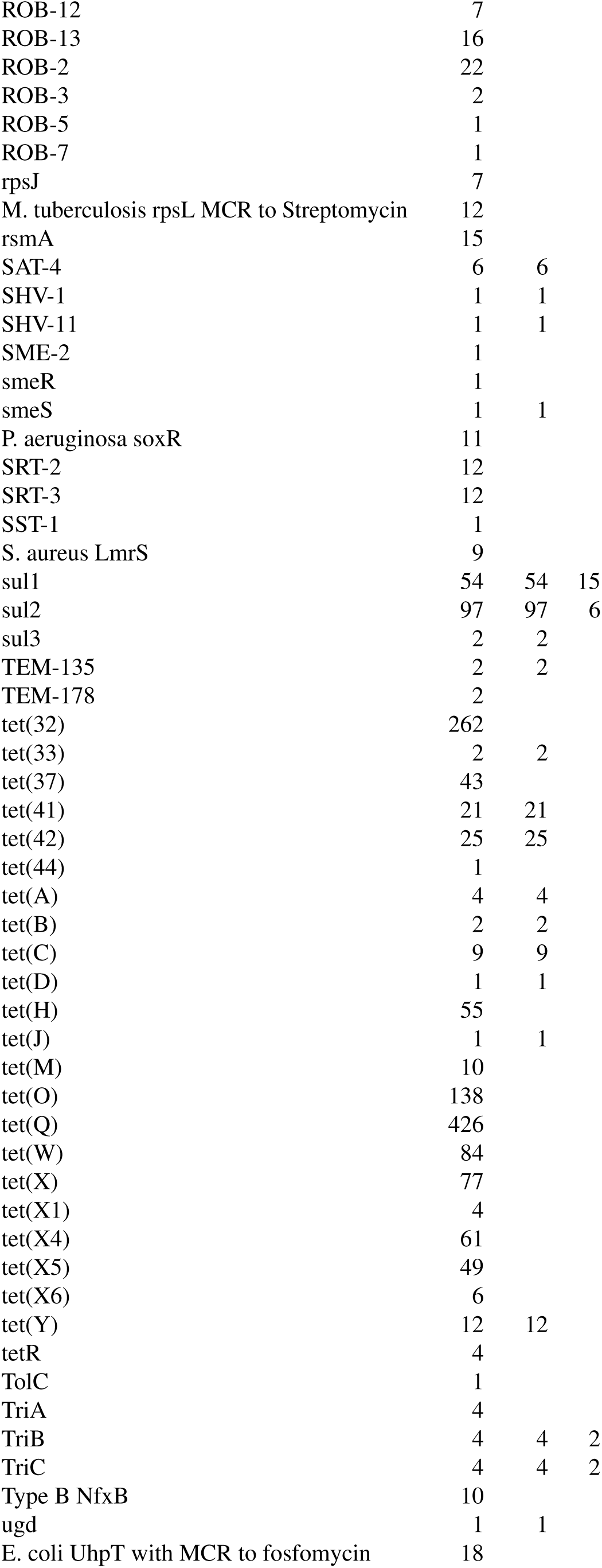
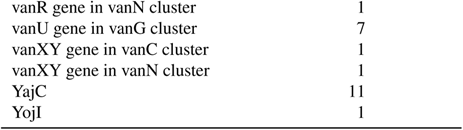
All ARG hits and the number of samples in which they have been detected (A), higher public health risk ARGs and the number of samples in which they have been detected (B) and higher public health risk ARGs deriving from ESKAPE pathogens and the number of samples in which they have been detected (C). MCR in the names is the abbreviation for mutation conferring resistance. *Escherichia coli* is abbreviated as E. coli, *Klebsiella pneumoniae* as K. pneumoniae, *Mycobacterium tuberculosis* as M. tuberculosis, Pseudomonas aeruginosa as P. aeruginosa and *Staphylococcus aureus* as S. aureus. The number of samples in which ARGs were detected is presented on axis X.

| ARG | A | B | C |
| --- | --- | --- | --- |
| 23S rRNA | 1 |  |  |
| (adenine(2058)-N(6))-methyltransferase |  |  |  |
| Erm(A) |  |  |  |
| AAC(3)-IId | 1 | 1 |  |
| AAC(3)-IV | 10 | 10 |  |
| AAC(6')-29b | 1 |  |  |
| AAC(6')-Iaa | 4 |  |  |
| AAC(6')-Iak | 4 |  |  |
| AAC(6')-Ib7 | 2 | 2 |  |
| AAC(6')-Ib9 | 2 | 2 |  |
| AAC(6')-Ic | 23 | 23 |  |
| AAC(6')-Im | 1 | 1 |  |
| AAC(6')-Iy | 4 |  |  |
| aad(6) | 1 | 1 |  |
| aadA2 | 13 | 13 |  |
| aadA22 | 1 | 1 |  |
| aadA25 | 1 |  |  |
| aadA27 | 1 |  |  |
| aadA3 | 6 | 6 |  |
| aadA6 | 1 | 1 |  |
| aadA8b | 1 |  |  |
| aadA9 | 2 |  |  |
| aadS | 119 |  |  |
| abeM | 1 | 1 | 1 |
| acrB | 13 | 13 | 4 |
| acrD | 11 | 11 | 2 |
| AcrE | 1 |  |  |
| AcrF | 1 |  |  |
| AcrS | 1 |  |  |
| ACT-2 | 1 |  |  |
| ACT-4 | 1 |  |  |
| ACT-56 | 1 |  |  |
| ADC-184 | 1 |  |  |
| adeF | 1 | 1 | 1 |
| adeG | 1 | 1 | 1 |
| adeH | 1 | 1 | 1 |
| adeI | 1 | 1 |  |
| adeK | 1 | 1 |  |
| almG | 4 |  |  |
| ANT(2'')-Ia | 19 | 19 | 3 |
| ANT(6)-Ia | 1 | 1 |  |
| ANT(6)-Ib | 5 | 5 |  |
| APH(2'')-IIa | 1 | 1 |  |
| APH(3'')-Ib | 130 | 130 | 7 |
| APH(3')-Ia | 54 | 54 | 5 |
| APH(3')-IIa | 2 | 2 |  |
| APH(3')-IIb | 8 | 8 | 6 |
| APH(3')-IIIa | 4 | 4 |  |
| APH(4)-Ia | 5 | 5 |  |
| APH(6)-Ic | 2 | 2 |  |
| APH(6)-Id | 153 | 153 | 17 |
| ArmR | 29 |  |  |
| arnA | 4 | 4 |  |
| ArnT | 4 |  |  |
| AxyX | 1 | 1 |  |
| bacA | 12 | 12 | 3 |
| baeR | 12 | 12 | 4 |
| baeS | 1 | 1 |  |
| basR | 7 |  |  |
| basS | 7 | 7 | 5 |
| bcr-1 | 6 | 6 | 4 |
| BLMT | 6 |  |  |
| BRO-2 | 1 |  |  |
| BRP(MBL) | 29 |  |  |
| CARB-16 | 3 | 3 |  |
| CARB-50 | 2 |  |  |
| catA4 | 1 |  |  |
| catB11 | 1 | 1 |  |
| catII | 13 | 13 |  |
| catIII | 14 | 14 | 2 |
| catS | 8 |  |  |
| catV | 28 |  |  |
| CfxA2 | 609 | 609 |  |
| CfxA3 | 44 | 44 |  |
| CfxA4 | 24 | 24 |  |
| CfxA5 | 7 | 7 |  |
| CfxA6 | 2 |  |  |
| chrB | 37 | 37 |  |
| cmlA1 | 2 | 2 |  |
| cmlA5 | 8 | 8 | 1 |
| cmlA9 | 2 |  |  |
| cmx | 2 | 2 |  |
| cprR | 9 |  |  |
| cprS | 6 |  |  |
| cpxA | 7 | 7 |  |
| CRP | 75 | 75 | 3 |
| CRP-1 | 1 |  |  |
| dfrA12 | 1 | 1 |  |
| dfrA14 | 12 | 12 | 4 |
| dfrA15 | 2 | 2 |  |
| dfrA20 | 30 |  |  |
| dfrC | 1 | 1 |  |
| dfrF | 1 | 1 |  |
| DHA-12 | 1 |  |  |
| DHA-14 | 1 |  |  |
| EC-15 | 1 |  |  |
| E. coli EF-Tu MCR to Pulvomycin | 1002 |  |  |
| emrA | 1 | 1 |  |
| emrB | 6 | 6 | 2 |
| emrK | 1 | 1 |  |
| emrR | 16 | 16 | 4 |
| emrY | 1 | 1 |  |
| Enterobacter cloacae acrA | 3 |  |  |
| eptA | 1 | 1 |  |
| eptB | 2 |  |  |
| EreA | 1 | 1 |  |
| EreD | 4 |  |  |
| Erm(41) | 19 |  |  |
| ErmB | 21 | 21 | 2 |
| ErmF | 202 | 202 |  |
| ErmG | 4 |  |  |
| ErmX | 7 | 7 |  |
| E. coli acrA | 1 |  |  |
| E. coli AcrAB-TolC with AcrR MCR to<br>ciprofloxacin, tetracycline, and ceftazidime | 1 |  |  |
| E. coli AcrAB-TolC with MarR MCR to<br>ciprofloxacin and tetracycline | 4 |  |  |
| E. coli EF-Tu MCR to kirromycin | 1 |  |  |
| E. coli emrE | 1 |  |  |
| E. coli mdfA | 1 |  |  |
| E. coli soxR with MCR | 1 |  |  |
| E. coli soxS with MCR | 9 |  |  |
| evgA | 1 | 1 |  |
| evgS | 1 | 1 |  |
| facT | 4 | 4 |  |
| floR | 1 | 1 |  |
| FosA | 11 |  |  |
| FosA2 | 4 | 4 | 2 |
| fosA5 | 1 |  |  |
| FosA6 | 2 | 2 | 1 |
| FosK | 2 |  |  |
| gadX | 1 | 1 |  |
| E. coli GlpT with MCR to fosfomycin | 13 |  |  |
| GOB-2 | 1 |  |  |
| H-NS | 12 | 12 | 2 |
| IND-6 | 1 |  |  |
| kdpE | 1 | 1 |  |
| K. pneumoniae acrA | 2 |  |  |
| K. pneumoniae KpnE | 4 |  |  |
| K. pneumoniae KpnF | 5 |  |  |
| K. pneumoniae KpnG | 5 |  |  |
| K. pneumoniae KpnH | 10 |  |  |
| K. pneumoniae OmpK37 | 3 |  |  |
| LEN-14 | 1 | 1 | 1 |
| lnuA | 1 | 1 | 1 |
| lnuB | 1 | 1 |  |
| lnuC | 4 | 4 |  |
| LpsB | 1 |  |  |
| LptD | 4 |  |  |
| lsaE | 1 | 1 |  |
| marA | 22 | 22 | 5 |
| MCR-9.1 | 1 |  |  |
| mdtA | 1 | 1 |  |
| mdtB | 9 | 9 | 1 |
| mdtC | 11 | 11 | 3 |
| mdtE | 1 | 1 |  |
| mdtF | 1 | 1 |  |
| mdtG | 1 | 1 |  |
| mdtH | 1 | 1 |  |
| mdtM | 1 | 1 |  |
| mdtN | 1 | 1 |  |
| mdtO | 1 | 1 |  |
| mdtP | 1 | 1 |  |
| MecC-type methicillin resistance repressor | 2 |  |  |
| MecI |  |  |  |
| mecI | 3 | 3 |  |
| Mef(En2) | 68 | 68 |  |
| mel | 18 | 18 |  |
| MexA | 8 | 8 | 2 |
| MexB | 5 | 5 |  |
| MexC | 7 | 7 | 5 |
| MexD | 4 | 4 | 2 |
| MexE | 5 | 5 | 3 |
| MexF | 5 | 5 | 3 |
| MexG | 12 |  |  |
| MexH | 9 |  |  |
| MexI | 3 |  |  |
| MexJ | 8 |  |  |
| MexK | 5 |  |  |
| MexL | 9 |  |  |
| mexM | 4 | 4 | 3 |
| mexN | 1 | 1 |  |
| mexP | 7 | 7 | 5 |
| mexQ | 5 | 5 | 3 |
| MexR | 9 |  |  |
| MexS | 6 |  |  |
| MexV | 6 |  |  |
| MexW | 6 |  |  |
| mexY | 5 | 5 | 5 |
| MexZ | 8 |  |  |
| Morganella morganii gyrB conferring<br>resistance to fluoroquinolones | 2 |  |  |
| mphA | 9 | 9 |  |
| mphE | 4 | 4 | 2 |
| msbA | 14 | 14 | 3 |
| msrE | 4 | 4 | 2 |
| mtrA | 1 | 1 |  |
| MuxA | 7 |  |  |
| MuxB | 5 |  |  |
| MuxC | 5 |  |  |
| M. tuberculosis intrinsic murA conferring<br>resistance to fosfomycin | 1 |  |  |
| nalC | 10 |  |  |
| nalD | 11 |  |  |
| nimA | 9 |  |  |
| nimC | 1 |  |  |
| nimF | 109 |  |  |
| nimJ | 1 |  |  |
| nimK | 1 |  |  |
| OmpA | 5 |  |  |
| OpmB | 6 | 6 | 4 |
| OpmD | 3 |  |  |
| opmE | 7 | 7 | 4 |
| OpmH | 9 | 9 | 5 |
| OprJ | 2 | 2 |  |
| OprM | 6 | 6 | 2 |
| OprN | 5 | 5 | 3 |
| oqxA | 8 | 8 | 4 |
| oqxB | 2 | 2 | 1 |
| OXA-114l | 1 |  |  |
| OXA-119 | 1 |  |  |
| OXA-2 | 104 | 104 | 18 |
| OXA-347 | 27 |  |  |
| OXA-494 | 3 |  |  |
| OXA-50 | 1 | 1 | 1 |
| OXA-540 | 1 |  |  |
| OXA-789 | 1 |  |  |
| OXA-838 | 4 |  |  |
| OXA-846 | 1 |  |  |
| OXA-851 | 1 |  |  |
| OXA-903 | 2 |  |  |
| OXA-904 | 1 |  |  |
| OXY-1-2 | 1 |  |  |
| ParR | 8 |  |  |
| ParS | 7 |  |  |
| patB | 1 | 1 |  |
| PDC-121 | 1 |  |  |
| PDC-23 | 1 |  |  |
| PDC-24 | 1 |  |  |
| PDC-3 | 2 | 2 | 1 |
| PDC-39 | 1 |  |  |
| PDC-46 | 1 |  |  |
| PDC-5 | 1 |  |  |
| PDC-98 | 1 |  |  |
| pgpB | 953 |  |  |
| PmpM | 5 | 5 | 3 |
| PmrF | 1 |  |  |
| P. aeruginosa catB7 | 9 |  |  |
| P. aeruginosa CpxR | 5 |  |  |
| P. aeruginosa emrE | 9 |  |  |
| P. aeruginosa parE conferring resistance to<br>fluoroquinolones | 1 |  |  |
| qacE | 2 |  |  |
| qacEdelta1 | 91 |  |  |
| qacG | 3 |  |  |
| qacJ | 1 |  |  |
| qacL | 59 |  |  |
| qnrE1 | 1 |  |  |
| QnrS1 | 2 | 2 |  |
| RAHN-1 | 1 |  |  |
| ramA | 9 | 9 | 2 |
| rmtD2 | 1 |  |  |
| ROB-1 | 474 |  |  |
| ROB-10 | 127 |  |  |
| ROB-11 | 24 |  |  |
| ROB-12 | 7 |  |  |
| ROB-13 | 16 |  |  |
| ROB-2 | 22 |  |  |
| ROB-3 | 2 |  |  |
| ROB-5 | 1 |  |  |
| ROB-7 | 1 |  |  |
| rpsJ | 7 |  |  |
| M. tuberculosis rpsL MCR to Streptomycin | 12 |  |  |
| rsmA | 15 |  |  |
| SAT-4 | 6 | 6 |  |
| SHV-1 | 1 | 1 |  |
| SHV-11 | 1 | 1 |  |
| SME-2 | 1 |  |  |
| smeR | 1 |  |  |
| smeS | 1 | 1 |  |
| P. aeruginosa soxR | 11 |  |  |
| SRT-2 | 12 |  |  |
| SRT-3 | 12 |  |  |
| SST-1 | 1 |  |  |
| S. aureus LmrS | 9 |  |  |
| sul1 | 54 | 54 | 15 |
| sul2 | 97 | 97 | 6 |
| sul3 | 2 | 2 |  |
| TEM-135 | 2 | 2 |  |
| TEM-178 | 2 |  |  |
| tet(32) | 262 |  |  |
| tet(33) | 2 | 2 |  |
| tet(37) | 43 |  |  |
| tet(41) | 21 | 21 |  |
| tet(42) | 25 | 25 |  |
| tet(44) | 1 |  |  |
| tet(A) | 4 | 4 |  |
| tet(B) | 2 | 2 |  |
| tet(C) | 9 | 9 |  |
| tet(D) | 1 | 1 |  |
| tet(H) | 55 |  |  |
| tet(J) | 1 | 1 |  |
| tet(M) | 10 |  |  |
| tet(O) | 138 |  |  |
| tet(Q) | 426 |  |  |
| tet(W) | 84 |  |  |
| tet(X) | 77 |  |  |
| tet(X1) | 4 |  |  |
| tet(X4) | 61 |  |  |
| tet(X5) | 49 |  |  |
| tet(X6) | 6 |  |  |
| tet(Y) | 12 | 12 |  |
| tetR | 4 |  |  |
| TolC | 1 |  |  |
| TriA | 4 |  |  |
| TriB | 4 | 4 | 2 |
| TriC | 4 | 4 | 2 |
| Type B NfxB | 10 |  |  |
| ugd | 1 | 1 |  |
| E. coli UhpT with MCR to fosfomycin | 18 |  |  |

|  |  |
| --- | --- |
| vanR gene in vanN cluster | 1 |
| vanU gene in vanG cluster | 7 |
| vanXY gene in vanC cluster | 1 |
| vanXY gene in vanN cluster | 1 |
| YajC | 11 |
| YojI | 1 |

**Table 6.**
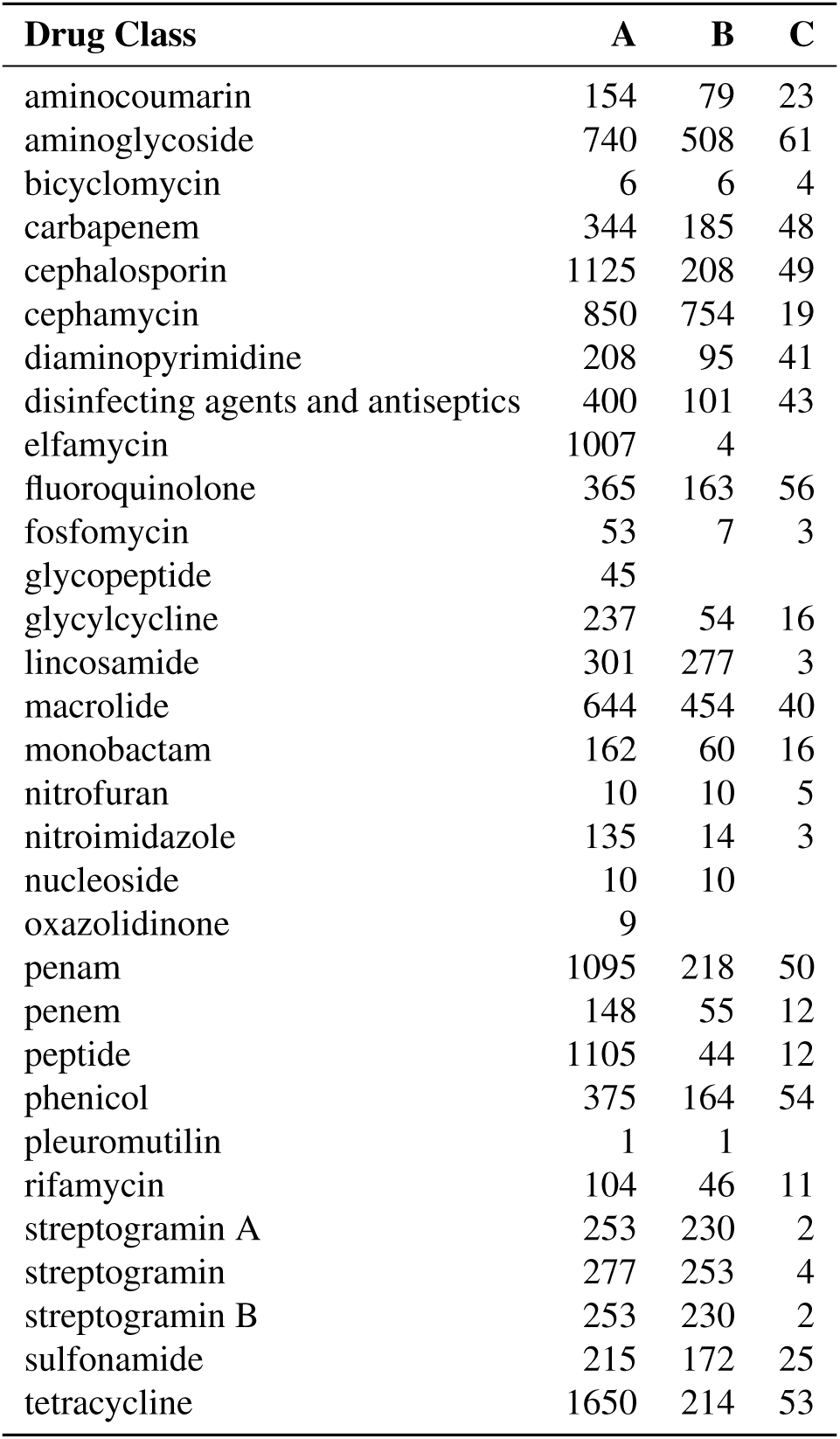
Antibiotic groups and the number of samples in whicg ARGs were detected against them. Antibiotic compounds affected by multidrug resistance are displayed separately.

**Table 7.** The number of samples containing ARGs (ARG+) and lacking ARGs (ARG-) associated with questions regarding the behaviour of dogs involved in the survey. ’Group 1’ and ’Group 2’ represent answer ’Never’ and ’Not never’ (Always, Often, Sometimes, Rarely), respectively. In column ’Approach’, ’A’ indicates ARGs detected in any canine metagenomic samples, ’B’ is for higher public health risk ARGs detected in any canine metagenomic samples and ’C’ stands for higher public health risk ARGs detected in ESKAPE pathogens.

| Question | Approach | Number of samples |  |  |  | OR (95%CI) | p-value |
| --- | --- | --- | --- | --- | --- | --- | --- |
|  |  | Never |  | Not never |  |  |  |
|  |  | AGR+ | AGR- | AGR+ | AGR- |  |  |
| Dog howls | A | 893 | 84 | 664 | 50 | 0.80 (0.54 - 1.17) | 0.238 |
|  | B | 546 | 432 | 402 | 312 | 0.98 (0.80 - 1.2)0 | 0.882 |
|  | C | 35 | 943 | 24 | 690 | 1.07 (0.61 - 1.89) | 0.894 |
| Dog woo-woo barks | A | 507 | 47 | 1055 | 84 | 0.86 (0.58 - 1.28) | 0.439 |
|  | B | 315 | 240 | 636 | 503 | 1.04 (0.84 - 1.28) | 0.754 |
|  | C | 15 | 540 | 45 | 1094 | 0.68 (0.35 - 1.25) | 0.210 |
| Dog whines when alone | A | 588 | 41 | 971 | 93 | 1.37 (0.93 - 2.06) | 0.113 |
|  | B | 357 | 272 | 592 | 473 | 1.05 (0.86 - 1.29) | 0.649 |
|  | C | 23 | 606 | 35 | 1030 | 1.12 (0.62 - 1.97) | 0.681 |
| Dog whines to get attention, food, or toys | A | 462 | 36 | 1047 | 94 | 1.15 (0.76 - 1.77) | 0.551 |
|  | B | 284 | 214 | 640 | 502 | 1.04 (0.84 - 1.30) | 0.745 |
|  | C | 19 | 479 | 40 | 1102 | 1.09 (0.59 - 1.95) | 0.774 |
| Dog eats grass | A | 162 | 13 | 1385 | 121 | 1.09 (0.60 - 2.15) | 0.883 |
|  | B | 108 | 68 | 835 | 671 | 1.28 (0.92 - 1.79) | 0.149 |
|  | C | 9 | 167 | 50 | 1456 | 1.57 (0.67 - 3.30) | 0.274 |
| Dog buries toys / bones | A | 1010 | 92 | 538 | 40 | 0.82 (0.54 - 1.22) | 0.340 |
|  | B | 621 | 482 | 320 | 258 | 1.04 (0.84 - 1.28) | 0.718 |
|  | C | 35 | 1068 | 23 | 555 | 0.79 (0.45 - 1.42) | 0.401 |
| Dog eats non-food items | A | 586 | 44 | 912 | 84 | 1.23 (0.83 - 1.84) | 0.300 |
|  | B | 364 | 266 | 553 | 444 | 1.10 (0.89 - 1.35) | 0.383 |
|  | C | 22 | 608 | 36 | 961 | 0.97 (0.54 - 1.71) | 1.000 |
| Dog escapes from cages, kennels, or enclosures | A | 852 | 71 | 694 | 61 | 1.05 (0.73 - 1.53) | 0.785 |
|  | B | 513 | 411 | 430 | 325 | 0.94 (0.77 - 1.15) | 0.587 |
|  | C | 37 | 887 | 23 | 732 | 1.33 (0.76 - 2.36) | 0.355 |
| Dog gets stuck behind objects and is unable to get around | A | 1048 | 93 | 498 | 39 | 0.88 (0.58 - 1.32) | 0.561 |
|  | B | 626 | 515 | 317 | 221 | 0.85 (0.68 - 1.05) | 0.126 |
|  | C | 35 | 1106 | 25 | 513 | 0.65 (0.37 - 1.15) | 0.121 |
| Dog walks away or avoids being patted | A | 705 | 50 | 841 | 80 | 1.34 (0.92 - 1.98) | 0.120 |
|  | B | 422 | 333 | 521 | 401 | 0.98 (0.80 - 1.19) | 0.805 |
|  | C | 36 | 719 | 24 | 898 | 1.87 (1.08 - 3.31) | 0.024* |
| Dog shows barrier aggression | A | 1037 | 82 | 507 | 50 | 1.25 (0.85 - 1.83) | 0.249 |
|  | B | 634 | 486 | 307 | 250 | 1.06 (0.86 - 1.31) | 0.566 |
|  | C | 43 | 1077 | 17 | 540 | 1.27 (0.70 - 2.39) | 0.486 |
| Dog lays in one place all day long | A | 734 | 68 | 815 | 64 | 0.85 (0.58 - 1.23) | 0.366 |
|  | B | 434 | 368 | 513 | 367 | 0.84 (0.69 - 1.03) | 0.085 |
|  | C | 28 | 774 | 31 | 849 | 0.99 (0.57 - 1.72) | 1.000 |
| Dog damages doors, gates, or walls | A | 1114 | 97 | 381 | 32 | 0.96 (0.62 - 1.48) | 0.916 |
|  | B | 687 | 525 | 228 | 185 | 1.06 (0.84 - 1.34) | 0.606 |
|  | C | 46 | 1166 | 13 | 400 | 1.21 (0.64 - 2.48) | 0.648 |
| Dog has difficulty finding food dropped on the floor | A | 751 | 62 | 794 | 68 | 1.04 (0.71 - 1.51) | 0.856 |
|  | B | 455 | 358 | 486 | 376 | 0.98 (0.81 - 1.20) | 0.883 |
|  | C | 25 | 788 | 34 | 828 | 0.77 (0.44 - 1.35) | 0.356 |
| Dog licks his empty bowl after finishing the food | A | 128 | 6 | 1410 | 124 | 1.88 (0.81 - 5.31) | 0.177 |
|  | B | 71 | 63 | 870 | 664 | 0.86 (0.59 - 1.25) | 0.415 |
|  | C | 5 | 129 | 54 | 1480 | 1.06 (0.33 - 2.70) | 0.808 |
| Dog licks tile / linoleum floors | A | 558 | 42 | 983 | 89 | 1.20 (0.81 - 1.81) | 0.393 |
|  | B | 330 | 270 | 608 | 464 | 0.93 (0.76 - 1.15) | 0.505 |
|  | C | 19 | 581 | 40 | 1032 | 0.84 (0.46 - 1.51) | 0.584 |
| Dog turns in circles before pooping | A | 383 | 34 | 1156 | 95 | 0.93 (0.61 - 1.44) | 0.751 |
|  | B | 241 | 176 | 696 | 555 | 1.09 (0.87 - 1.38) | 0.459 |
|  | C | 15 | 402 | 44 | 1207 | 1.02 (0.52 - 1.90) | 1.000 |
| Dog kicks or scratches the ground after defecating | A | 385 | 37 | 1153 | 91 | 0.82 (0.54 - 1.26) | 0.342 |
|  | B | 238 | 184 | 698 | 546 | 1.01 (0.81 - 1.27) | 0.955 |
|  | C | 11 | 411 | 48 | 1196 | 0.67 (0.31 - 1.32) | 0.286 |
| Dog lifts his leg to urinate | A | 598 | 46 | 935 | 84 | 1.17 (0.79 - 1.74) | 0.454 |
|  | B | 351 | 293 | 583 | 436 | 0.90 (0.73 - 1.10) | 0.287 |
|  | C | 21 | 623 | 36 | 983 | 0.92 (0.51 - 1.64) | 0.890 |
| Dog marks with feces | A | 1034 | 91 | 502 | 39 | 0.88 (0.58 - 1.32) | 0.560 |
|  | B | 624 | 501 | 311 | 230 | 0.92 (0.74 - 1.14) | 0.461 |
|  | C | 42 | 1083 | 16 | 525 | 1.27 (0.69 - 2.45) | 0.477 |
| Dog eats his own feces | A | 1310 | 108 | 230 | 21 | 1.11 (0.65 - 1.82) | 0.700 |
|  | B | 795 | 623 | 143 | 108 | 0.96 (0.73 - 1.27) | 0.836 |
|  | C | 50 | 1368 | 9 | 242 | 0.98 (0.47 - 2.30) | 1.000 |
| Dog shows submissive urination | A | 1170 | 100 | 370 | 31 | 0.98 (0.62 - 1.51) | 1.000 |
|  | B | 726 | 544 | 214 | 187 | 1.17 (0.92 - 1.47) | 0.185 |
|  | C | 46 | 1224 | 13 | 388 | 1.12 (0.59 - 2.29) | 0.877 |
| Dog drinks water quickly | A | 117 | 5 | 1371 | 122 | 2.08 (0.84 - 6.66) | 0.117 |
|  | B | 76 | 46 | 833 | 660 | 1.31 (0.88 - 1.96) | 0.184 |
|  | C | 4 | 118 | 54 | 1439 | 0.90 (0.23 - 2.51) | 1.000 |
| Dog rests frog style | A | 512 | 37 | 1015 | 92 | 1.25 (0.83 - 1.92) | 0.285 |
|  | B | 309 | 240 | 621 | 486 | 1.01 (0.82 - 1.25) | 0.958 |
|  | C | 19 | 530 | 38 | 1069 | 1.01 (0.54 - 1.81) | 1.000 |
| Dog stares blankly at the walls or floor | A | 909 | 83 | 610 | 47 | 0.84 (0.57 - 1.24) | 0.402 |
|  | B | 553 | 439 | 374 | 283 | 0.95 (0.78 - 1.17) | 0.648 |
|  | C | 34 | 958 | 24 | 633 | 0.94 (0.53 - 1.67) | 0.892 |
| Dog tilts his head | A | 140 | 20 | 1384 | 110 | 0.56 (0.33 - 0.98) | 0.029* |
|  | B | 92 | 68 | 838 | 656 | 1.06 (0.75 - 1.50) | 0.801 |
|  | C | 5 | 155 | 53 | 1441 | 0.88 (0.27 - 2.23) | 1.000 |
| Dog crosses his front paws | A | 338 | 29 | 1185 | 100 | 0.98 (0.63 - 1.57) | 0.913 |
|  | B | 199 | 168 | 728 | 557 | 0.91 (0.71 - 1.15) | 0.438 |
|  | C | 10 | 357 | 48 | 1237 | 0.72 (0.32 - 1.46) | 0.423 |
| Dog shows handedness or side-preference | A | 381 | 42 | 1140 | 87 | 0.69 (0.46 - 1.05) | 0.074 |
|  | B | 240 | 183 | 688 | 539 | 1.03 (0.82 - 1.29) | 0.820 |
|  | C | 12 | 411 | 45 | 1182 | 0.77 (0.37 - 1.49) | 0.537 |
| Dog places his paw on my or other people's feet | A | 295 | 29 | 1177 | 98 | 0.85 (0.54 - 1.36) | 0.490 |
|  | B | 190 | 134 | 711 | 564 | 1.12 (0.87 - 1.45) | 0.380 |
|  | C | 10 | 314 | 47 | 1228 | (0.83 0.37 - 1.69) | 0.738 |

**Table 8.**
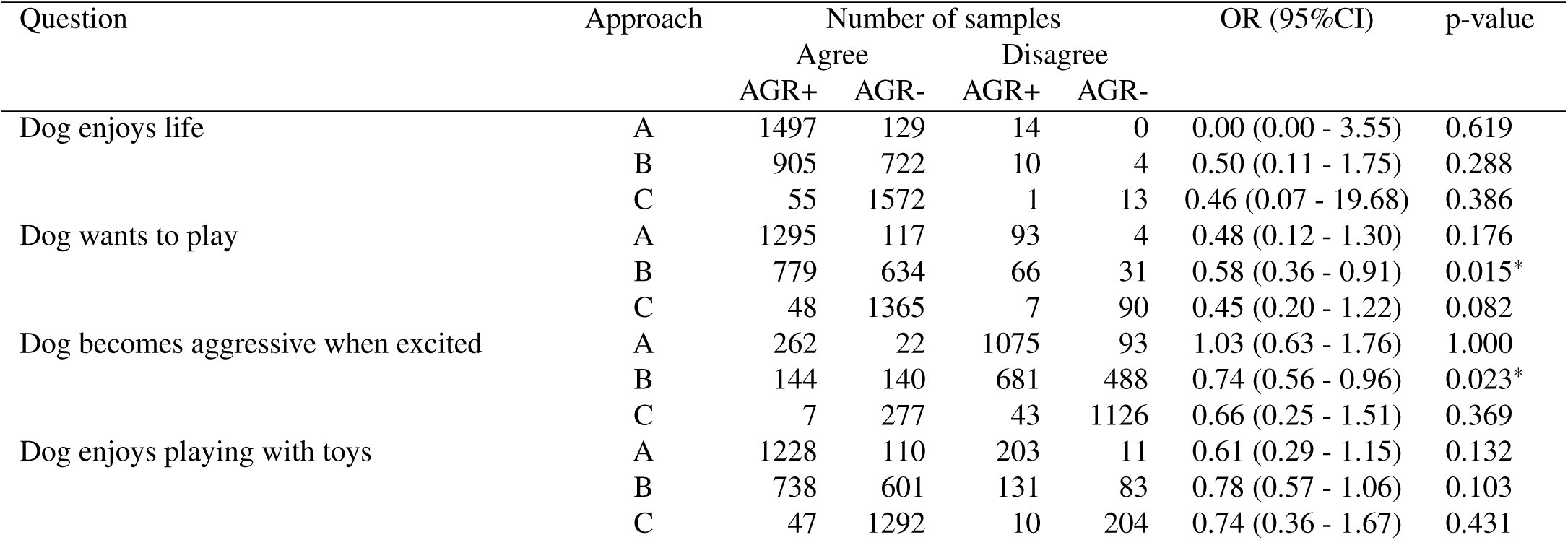
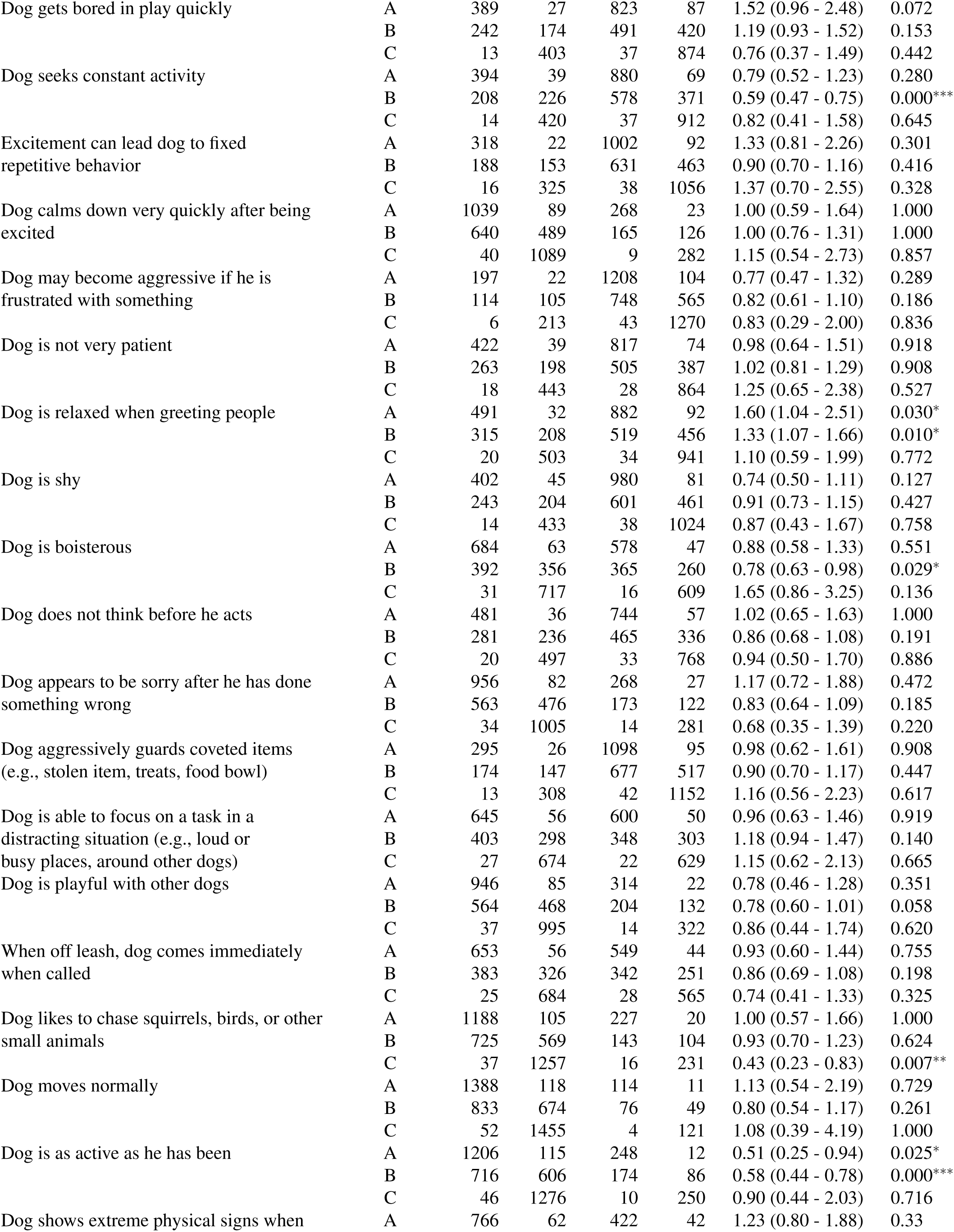
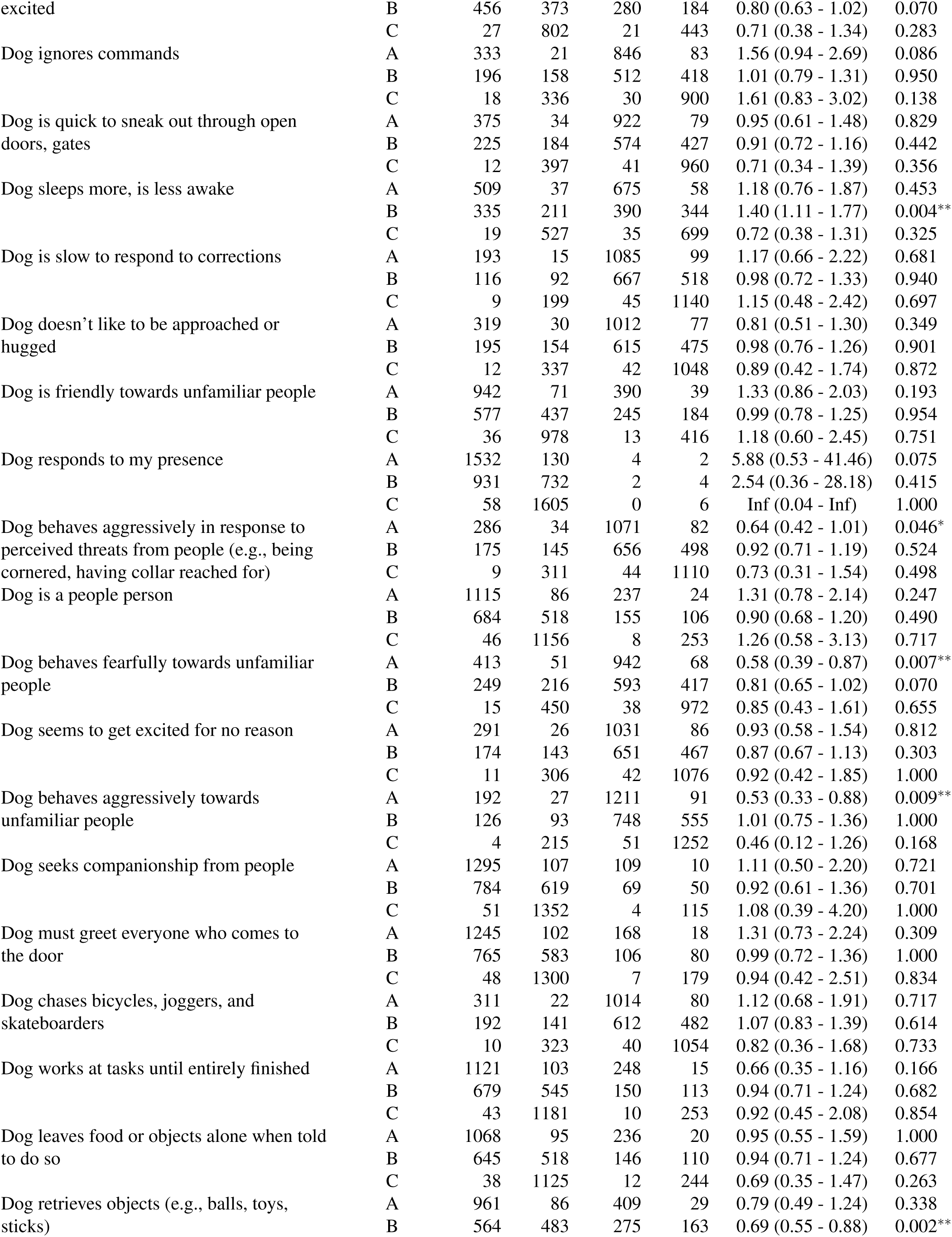
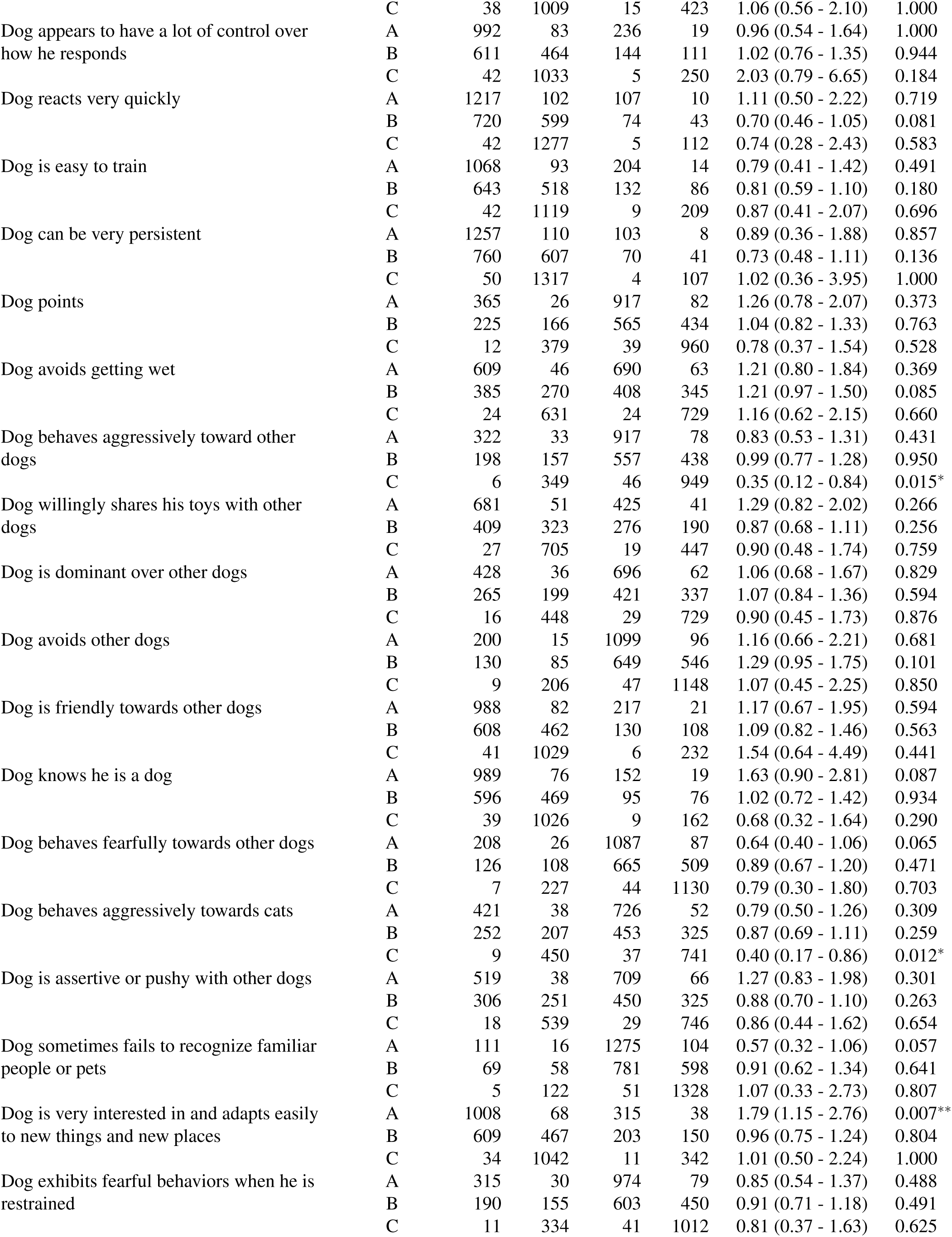
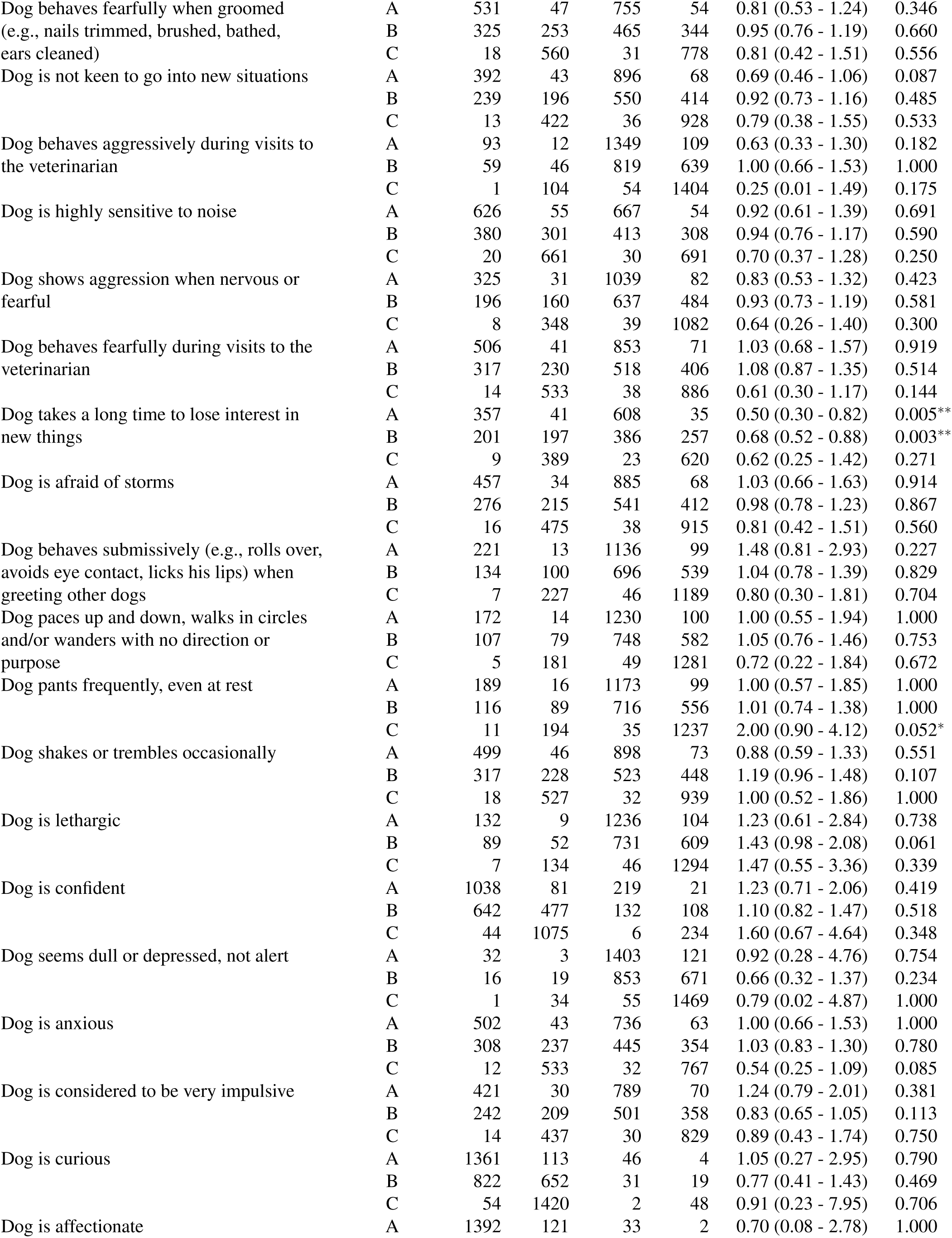
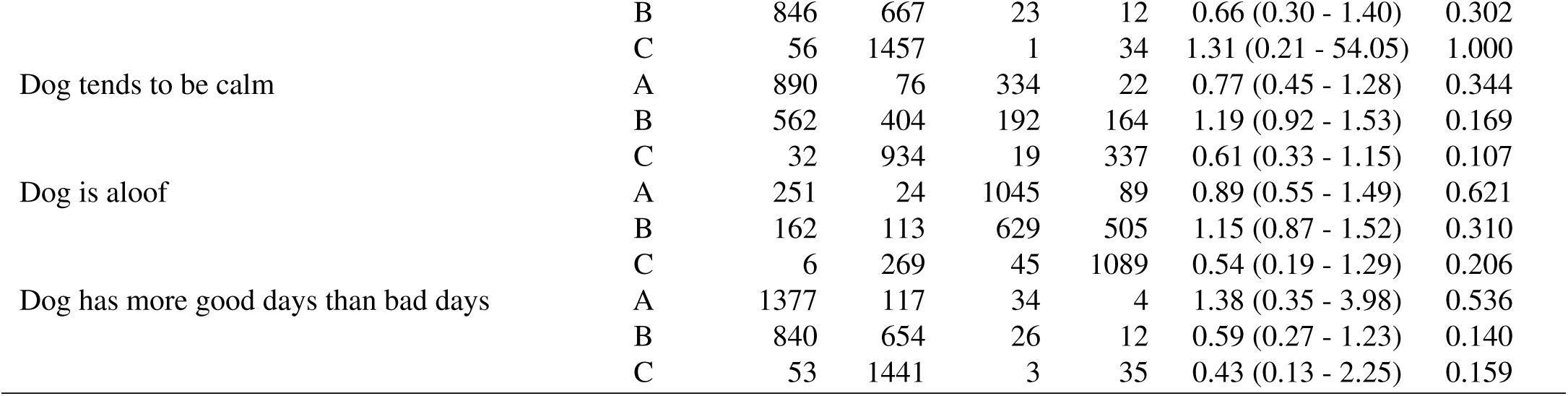
The number of samples containing ARGs (ARG+) and lacking ARGs (ARG-) associated with questions regarding the behaviour of dogs involved in the survey. ’Group 1’ and ’Group 2’ represent answer ’Agree’ (Strongly agree, Agree) and ’Disagree’ (Strongly disagree, disagree), respectively. In column ’Approach’, ’A’ indicates ARGs detected in any canine metagenomic samples, ’B’ is for higher public health risk ARGs detected in any canine metagenomic samples and ’C’ stands for higher public health risk ARGs detected in ESKAPE pathogens.

**Table 9.** The number of samples containing ARGs (ARG+) and lacking ARGs (ARG-) associated with questions regarding the behaviour of dogs involved in the survey. ’Group 1’ and ’Group 2’ represent answer ’Agree’ (Strongly agree, Agree), ’Neither agree, nor disagree’ and ’Disagree’ (Strongly disagree, disagree), respectively. In column ’Approach’, ’A’ indicates ARGs detected in any canine metagenomic samples, ’B’ is for higher public health risk ARGs detected in any canine metagenomic samples and ’C’ stands for higher public health risk ARGs detected in ESKAPE pathogens.

| Question | Approach | Number of samples |  |  |  |  |  | p-value |
| --- | --- | --- | --- | --- | --- | --- | --- | --- |
|  |  | Agree |  | Neither |  | Disagree |  |  |
|  |  | ARG+ | ARG- | ARG+ | ARG- | ARG+ | ARG- |  |
| Dog enjoys life | A | 1497 | 129 | 60 | 4 | 14 | 0 | 0.741 |
|  | B | 905 | 722 | 43 | 21 | 10 | 4 | 0.095 |
|  | C | 55 | 1572 | 3 | 61 | 1 | 13 | 0.332 |
| Dog wants to play | A | 1295 | 117 | 177 | 10 | 93 | 4 | 0.167 |
|  | B | 779 | 634 | 109 | 78 | 66 | 31 | 0.037 |
|  | C | 48 | 1365 | 4 | 183 | 7 | 90 | 0.097 |
| Dog becomes aggressive when excited | A | 262 | 22 | 226 | 16 | 1075 | 93 | 0.815 |
|  | B | 144 | 140 | 125 | 117 | 681 | 488 | 0.024* |
|  | C | 7 | 277 | 8 | 234 | 43 | 1126 | 0.644 |
| Dog enjoys playing with toys | A | 1228 | 110 | 140 | 13 | 203 | 11 | 0.283 |
|  | B | 738 | 601 | 88 | 65 | 131 | 83 | 0.233 |
|  | C | 47 | 1292 | 2 | 151 | 10 | 204 | 0.216 |
| Dog gets bored in play quickly | A | 389 | 27 | 349 | 16 | 823 | 87 | 0.004** |
|  | B | 242 | 174 | 218 | 147 | 491 | 420 | 0.108 |
|  | C | 13 | 403 | 9 | 356 | 37 | 874 | 0.348 |
| Dog seeks constant activity | A | 394 | 39 | 288 | 26 | 880 | 69 | 0.499 |
|  | B | 208 | 226 | 166 | 148 | 578 | 371 | 0.000*** |
|  | C | 14 | 420 | 7 | 307 | 37 | 912 | 0.390 |
| Excitement can lead DOG to fixed repetitive behavior | A | 318 | 22 | 242 | 19 | 1002 | 92 | 0.503 |
|  | B | 188 | 153 | 135 | 126 | 631 | 463 | 0.198 |
|  | C | 16 | 325 | 5 | 256 | 38 | 1056 | 0.181 |
| Dog calms down very quickly after being excited | A | 1039 | 89 | 253 | 20 | 268 | 23 | 0.972 |
|  | B | 640 | 489 | 145 | 128 | 165 | 126 | 0.557 |
|  | C | 40 | 1089 | 10 | 263 | 9 | 282 | 0.922 |
| Dog may become aggressive if he is frustrated with something | A | 197 | 22 | 156 | 8 | 1208 | 104 | 0.174 |
|  | B | 114 | 105 | 91 | 73 | 748 | 565 | 0.384 |
|  | C | 6 | 213 | 10 | 154 | 43 | 1270 | 0.164 |
| Dog is not very patient | A | 422 | 39 | 275 | 18 | 817 | 74 | 0.456 |
|  | B | 263 | 198 | 162 | 131 | 505 | 387 | 0.892 |
|  | C | 18 | 443 | 12 | 281 | 28 | 864 | 0.618 |
| Dog is relaxed when greeting people | A | 491 | 32 | 193 | 10 | 882 | 92 | 0.019* |
|  | B | 315 | 208 | 120 | 83 | 519 | 456 | 0.022* |
|  | C | 20 | 503 | 6 | 197 | 34 | 941 | 0.877 |
| Dog is shy | A | 402 | 45 | 171 | 8 | 980 | 81 | 0.053* |
|  | B | 243 | 204 | 102 | 77 | 601 | 461 | 0.702 |
|  | C | 14 | 433 | 8 | 171 | 38 | 1024 | 0.677 |
| Dog is boisterous | A | 684 | 63 | 298 | 24 | 578 | 47 | 0.786 |
|  | B | 392 | 356 | 192 | 130 | 365 | 260 | 0.029* |
|  | C | 31 | 717 | 13 | 309 | 16 | 609 | 0.233 |
| Dog does not think before he acts | A | 481 | 36 | 338 | 41 | 744 | 57 | 0.066 |
|  | B | 281 | 236 | 205 | 175 | 465 | 336 | 0.274 |
|  | C | 20 | 497 | 7 | 373 | 33 | 768 | 0.112 |
| Dog appears to be sorry after he has done something wrong | A | 956 | 82 | 333 | 25 | 268 | 27 | 0.594 |
|  | B | 563 | 476 | 213 | 145 | 173 | 122 | 0.137 |
|  | C | 34 | 1005 | 10 | 348 | 14 | 281 | 0.363 |
| Dog aggressively guards coveted items (e.g., stolen item, treats, food bowl) | A | 295 | 26 | 165 | 13 | 1098 | 95 | 0.968 |
|  | B | 174 | 147 | 99 | 79 | 677 | 517 | 0.714 |
|  | C | 13 | 308 | 5 | 173 | 42 | 1152 | 0.793 |
| Dog is able to focus on a task in a distracting situation (e.g., loud or busy places, around other dogs) | A | 645 | 56 | 313 | 27 | 600 | 50 | 0.979 |
|  | B | 403 | 298 | 198 | 142 | 348 | 303 | 0.221 |
|  | C | 27 | 674 | 10 | 330 | 22 | 629 | 0.769 |
| Dog is playful with other dogs | A | 946 | 85 | 289 | 25 | 314 | 22 | 0.633 |
|  | B | 564 | 468 | 174 | 140 | 204 | 132 | 0.146 |
|  | C | 37 | 995 | 7 | 307 | 14 | 322 | 0.370 |
| When off leash, dog comes immediately when called | A | 653 | 56 | 345 | 30 | 549 | 44 | 0.930 |
|  | B | 383 | 326 | 221 | 155 | 342 | 251 | 0.235 |
|  | C | 25 | 684 | 6 | 370 | 28 | 565 | 0.030 |
| Dog likes to chase squirrels, birds, or other small animals | A | 1188 | 105 | 133 | 9 | 227 | 20 | 0.821 |
|  | B | 725 | 569 | 73 | 69 | 143 | 104 | 0.458 |
|  | C | 37 | 1257 | 5 | 137 | 16 | 231 | 0.024 |
| Dog moves normally | A | 1388 | 118 | 54 | 5 | 114 | 11 | 0.851 |
|  | B | 833 | 674 | 38 | 21 | 76 | 49 | 0.208 |
|  | C | 52 | 1455 | 3 | 56 | 4 | 121 | 0.698 |
| Dog is as active as he has been | A | 1206 | 115 | 89 | 5 | 248 | 12 | 0.052* |
|  | B | 716 | 606 | 51 | 43 | 174 | 86 | 0.001** |
|  | C | 46 | 1276 | 3 | 91 | 10 | 250 | 0.935 |
| Dog shows extreme physical signs when excited | A | 766 | 62 | 358 | 29 | 422 | 42 | 0.566 |
|  | B | 456 | 373 | 205 | 182 | 280 | 184 | 0.069 |
|  | C | 27 | 802 | 11 | 376 | 21 | 443 | 0.372 |
| Dog ignores commands | A | 333 | 21 | 377 | 28 | 846 | 83 | 0.157 |
|  | B | 196 | 158 | 240 | 165 | 512 | 418 | 0.343 |
|  | C | 18 | 336 | 12 | 393 | 30 | 900 | 0.219 |
| Dog is quick to sneak out through open doors, gates | A | 375 | 34 | 249 | 19 | 922 | 79 | 0.857 |
|  | B | 225 | 184 | 144 | 125 | 574 | 427 | 0.459 |
|  | C | 12 | 397 | 7 | 262 | 41 | 960 | 0.430 |
| Dog sleeps more, is less awake | A | 509 | 37 | 361 | 34 | 675 | 58 | 0.551 |
|  | B | 335 | 211 | 217 | 178 | 390 | 344 | 0.011* |
|  | C | 19 | 527 | 6 | 389 | 35 | 699 | 0.015* |
| Dog is slow to respond to corrections | A | 193 | 15 | 269 | 16 | 1085 | 99 | 0.295 |
|  | B | 116 | 92 | 163 | 122 | 667 | 518 | 0.948 |
|  | C | 9 | 199 | 6 | 279 | 45 | 1140 | 0.294 |
| Dog doesn't like to be approached or hugged | A | 319 | 30 | 224 | 24 | 1012 | 77 | 0.290 |
|  | B | 195 | 154 | 137 | 111 | 615 | 475 | 0.942 |
|  | C | 12 | 337 | 5 | 243 | 42 | 1048 | 0.385 |
| Dog is friendly towards unfamiliar people | A | 942 | 71 | 223 | 22 | 390 | 39 | 0.295 |
|  | B | 577 | 437 | 126 | 119 | 245 | 184 | 0.270 |
|  | C | 36 | 978 | 10 | 235 | 13 | 416 | 0.756 |
| Dog responds to my presence | A | 1532 | 130 | 18 | 0 | 4 | 2 | 0.054* |
|  | B | 931 | 732 | 14 | 4 | 2 | 4 | 0.097 |
|  | C | 58 | 1605 | 1 | 17 | 0 | 6 | 0.577 |
| Dog behaves aggressively in response to perceived threats from people | A | 286 | 34 | 195 | 14 | 1071 | 82 | 0.109 |
|  | B | 175 | 145 | 114 | 95 | 656 | 498 | 0.692 |
|  | C | 9 | 311 | 5 | 204 | 44 | 1110 | 0.547 |
| Dog is a people person | A | 1115 | 86 | 196 | 22 | 237 | 24 | 0.205 |
|  | B | 684 | 518 | 107 | 111 | 155 | 106 | 0.056 |
|  | C | 46 | 1156 | 4 | 214 | 8 | 253 | 0.371 |
| Dog behaves fearfully towards unfamiliar people | A | 413 | 51 | 196 | 13 | 942 | 68 | 0.015* |
|  | B | 249 | 216 | 104 | 105 | 593 | 417 | 0.025* |
|  | C | 15 | 450 | 5 | 204 | 38 | 972 | 0.624 |
| Dog seems to get excited for no reason | A | 291 | 26 | 228 | 18 | 1031 | 86 | 0.925 |
|  | B | 174 | 143 | 119 | 127 | 651 | 467 | 0.017* |
|  | C | 11 | 306 | 6 | 240 | 42 | 1076 | 0.690 |
| Dog behaves aggressively towards unfamiliar people | A | 192 | 27 | 146 | 14 | 1211 | 91 | 0.026* |
|  | B | 126 | 93 | 72 | 88 | 748 | 555 | 0.011* |
|  | C | 4 | 215 | 4 | 156 | 51 | 1252 | 0.291 |
| DOG seeks companionship from people | A | 1295 | 107 | 150 | 14 | 109 | 10 | 0.826 |
|  | B | 784 | 619 | 96 | 68 | 69 | 50 | 0.769 |
|  | C | 51 | 1352 | 4 | 160 | 4 | 115 | 0.832 |
| Dog must greet everyone who comes to the door | A | 1245 | 102 | 82 | 9 | 168 | 18 | 0.406 |
|  | B | 765 | 583 | 45 | 46 | 106 | 80 | 0.394 |
|  | C | 48 | 1300 | 2 | 89 | 7 | 179 | 0.854 |
| Dog chases bicycles, joggers, and skateboarders | A | 311 | 22 | 225 | 29 | 1014 | 80 | 0.070 |
|  | B | 192 | 141 | 136 | 118 | 612 | 482 | 0.612 |
|  | C | 10 | 323 | 9 | 245 | 40 | 1054 | 0.906 |
| Dog works at tasks (e.g., getting treats out of a Kong, shredding toys) until entirely finished | A | 1121 | 103 | 171 | 10 | 248 | 15 | 0.192 |
|  | B | 679 | 545 | 106 | 75 | 150 | 113 | 0.706 |
|  | C | 43 | 1181 | 5 | 176 | 10 | 253 | 0.860 |
| Dog leaves food or objects alone when told to do so | A | 1068 | 95 | 242 | 16 | 236 | 20 | 0.610 |
|  | B | 645 | 518 | 148 | 110 | 146 | 110 | 0.803 |
|  | C | 38 | 1125 | 9 | 249 | 12 | 244 | 0.487 |
| Dog retrieves objects (e.g., balls, toys, sticks) | A | 961 | 86 | 177 | 15 | 409 | 29 | 0.610 |
|  | B | 564 | 483 | 100 | 92 | 275 | 163 | 0.0033* |
|  | C | 38 | 1009 | 5 | 187 | 15 | 423 | 0.866 |
| Dog appears to have a lot of control over how he responds | A | 992 | 83 | 315 | 29 | 236 | 19 | 0.879 |
|  | B | 611 | 464 | 180 | 164 | 144 | 111 | 0.332 |
|  | C | 42 | 1033 | 12 | 332 | 5 | 250 | 0.346 |
| Dog reacts very quickly | A | 1217 | 102 | 223 | 19 | 107 | 10 | 0.889 |
|  | B | 720 | 599 | 145 | 97 | 74 | 43 | 0.080 |
|  | C | 42 | 1277 | 12 | 230 | 5 | 112 | 0.328 |
| Dog is easy to train | A | 1068 | 93 | 271 | 24 | 204 | 14 | 0.732 |
|  | B | 643 | 518 | 162 | 133 | 132 | 86 | 0.343 |
|  | C | 42 | 1119 | 8 | 287 | 9 | 209 | 0.686 |
| Dog can be very persistent | A | 1257 | 110 | 186 | 13 | 103 | 8 | 0.769 |
|  | B | 760 | 607 | 108 | 91 | 70 | 41 | 0.278 |
|  | C | 50 | 1317 | 4 | 195 | 4 | 107 | 0.525 |
| Dog points | A | 365 | 26 | 258 | 23 | 917 | 82 | 0.630 |
|  | B | 225 | 166 | 144 | 137 | 565 | 434 | 0.215 |
|  | C | 12 | 379 | 8 | 273 | 39 | 960 | 0.659 |
| Dog avoids getting wet | A | 609 | 46 | 189 | 19 | 690 | 63 | 0.488 |
|  | B | 385 | 270 | 114 | 94 | 408 | 345 | 0.204 |
|  | C | 24 | 631 | 8 | 200 | 24 | 729 | 0.798 |
| Dog behaves aggressively toward other dogs | A | 322 | 33 | 297 | 18 | 917 | 78 | 0.227 |
|  | B | 198 | 157 | 181 | 134 | 557 | 438 | 0.887 |
|  | C | 6 | 349 | 7 | 308 | 46 | 949 | 0.014 |
| Dog willingly shares his toys with other dogs | A | 681 | 51 | 418 | 36 | 425 | 41 | 0.502 |
|  | B | 409 | 323 | 245 | 209 | 276 | 190 | 0.260 |
|  | C | 27 | 705 | 12 | 442 | 19 | 447 | 0.459 |
| Dog is dominant over other dogs | A | 428 | 36 | 405 | 31 | 696 | 62 | 0.816 |
|  | B | 265 | 199 | 248 | 188 | 421 | 337 | 0.842 |
|  | C | 16 | 448 | 14 | 422 | 29 | 729 | 0.873 |
| Dog avoids other dogs | A | 200 | 15 | 237 | 18 | 1099 | 96 | 0.855 |
|  | B | 130 | 85 | 156 | 99 | 649 | 546 | 0.053* |
|  | C | 9 | 206 | 3 | 252 | 47 | 1148 | 0.063 |
| Dog is friendly towards other dogs | A | 988 | 82 | 325 | 24 | 217 | 21 | 0.673 |
|  | B | 608 | 462 | 196 | 153 | 130 | 108 | 0.823 |
|  | C | 41 | 1029 | 12 | 337 | 6 | 232 | 0.685 |
| Dog knows he is a dog | A | 989 | 76 | 389 | 33 | 152 | 19 | 0.199 |
|  | B | 596 | 469 | 241 | 181 | 95 | 76 | 0.909 |
|  | C | 39 | 1026 | 10 | 412 | 9 | 162 | 0.185 |
| Dog behaves fearfully towards other dogs | A | 208 | 26 | 237 | 14 | 1087 | 87 | 0.069 |
|  | B | 126 | 108 | 143 | 108 | 665 | 509 | 0.713 |
|  | C | 7 | 227 | 8 | 243 | 44 | 1130 | 0.873 |
| Dog behaves aggressively towards cats | A | 421 | 38 | 385 | 37 | 726 | 52 | 0.342 |
|  | B | 252 | 207 | 230 | 192 | 453 | 325 | 0.351 |
|  | C | 9 | 450 | 13 | 409 | 37 | 741 | 0.032 |
| Dog is assertive or pushy with other dogs (e.g., if in a home with other dogs, when greeting) | A | 519 | 38 | 302 | 24 | 709 | 66 | 0.526 |
|  | B | 306 | 251 | 177 | 149 | 450 | 325 | 0.380 |
|  | C | 18 | 539 | 11 | 315 | 29 | 746 | 0.894 |
| Dog sometimes fails to recognize familiar people or pets | A | 111 | 16 | 91 | 4 | 1275 | 104 | 0.062 |
|  | B | 69 | 58 | 56 | 39 | 781 | 598 | 0.791 |
|  | C | 5 | 122 | 2 | 93 | 51 | 1328 | 0.766 |
| Dog is very interested in and adapts easily to new things and new places | A | 1008 | 68 | 209 | 24 | 315 | 38 | 0.008** |
|  | B | 609 | 467 | 123 | 110 | 203 | 150 | 0.494 |
|  | C | 34 | 1042 | 14 | 219 | 11 | 342 | 0.110 |
| Dog exhibits fearful behaviors when he is restrained | A | 315 | 30 | 236 | 18 | 974 | 79 | 0.712 |
|  | B | 190 | 155 | 137 | 117 | 603 | 450 | 0.554 |
|  | C | 11 | 334 | 7 | 247 | 41 | 1012 | 0.678 |
| Dog behaves fearfully when groomed (e.g., nails trimmed, brushed, bathed, ears cleaned) | A | 531 | 47 | 244 | 28 | 755 | 54 | 0.145 |
|  | B | 325 | 253 | 141 | 131 | 465 | 344 | 0.270 |
|  | C | 18 | 560 | 9 | 263 | 31 | 778 | 0.776 |
| Dog is not keen to go into new situations | A | 392 | 43 | 236 | 16 | 896 | 68 | 0.138 |
|  | B | 239 | 196 | 144 | 108 | 550 | 414 | 0.745 |
|  | C | 13 | 422 | 10 | 242 | 36 | 928 | 0.735 |
| Dog behaves aggressively during visits to the veterinarian | A | 93 | 12 | 77 | 9 | 1349 | 109 | 0.190 |
|  | B | 59 | 46 | 47 | 39 | 819 | 639 | 0.958 |
|  | C | 1 | 104 | 2 | 84 | 54 | 1404 | 0.319 |
| Dog is highly sensitive to noise | A | 626 | 55 | 234 | 21 | 667 | 54 | 0.880 |
|  | B | 380 | 301 | 140 | 115 | 413 | 308 | 0.760 |
|  | C | 20 | 661 | 9 | 246 | 30 | 691 | 0.478 |
| Dog shows aggression when nervous or fearful | A | 325 | 31 | 159 | 17 | 1039 | 82 | 0.413 |
|  | B | 196 | 160 | 96 | 80 | 637 | 484 | 0.747 |
|  | C | 8 | 348 | 10 | 166 | 39 | 1082 | 0.136 |
| Dog behaves fearfully during visits to the veterinarian | A | 506 | 41 | 168 | 17 | 853 | 71 | 0.716 |
|  | B | 317 | 230 | 95 | 90 | 518 | 406 | 0.296 |
|  | C | 14 | 533 | 6 | 179 | 38 | 886 | 0.287 |
| Dog takes a long time to lose interest in new things | A | 357 | 41 | 564 | 54 | 608 | 35 | 0.009** |
|  | B | 201 | 197 | 346 | 272 | 386 | 257 | 0.011* |
|  | C | 9 | 389 | 26 | 592 | 23 | 620 | 0.253 |
| Dog is afraid of storms | A | 457 | 34 | 139 | 25 | 885 | 68 | 0.003** |
|  | B | 276 | 215 | 89 | 75 | 541 | 412 | 0.833 |
|  | C | 16 | 475 | 4 | 160 | 38 | 915 | 0.631 |
| Dog behaves submissively when greeting other dogs | A | 221 | 13 | 173 | 18 | 1136 | 99 | 0.287 |
|  | B | 134 | 100 | 103 | 88 | 696 | 539 | 0.764 |
|  | C | 7 | 227 | 5 | 186 | 46 | 1189 | 0.774 |
| Dog paces up and down, walks in circles, etc. | A | 172 | 14 | 116 | 15 | 1230 | 100 | 0.275 |
|  | B | 107 | 79 | 71 | 60 | 748 | 582 | 0.844 |
|  | C | 5 | 181 | 2 | 129 | 49 | 1281 | 0.466 |
| Dog pants frequently, even at rest | A | 189 | 16 | 155 | 14 | 1173 | 99 | 0.961 |
|  | B | 116 | 89 | 96 | 73 | 716 | 556 | 0.992 |
|  | C | 11 | 194 | 12 | 157 | 35 | 1237 | 0.006 |
| Dog shakes or trembles occasionally | A | 499 | 46 | 124 | 10 | 898 | 73 | 0.809 |
|  | B | 317 | 228 | 86 | 48 | 523 | 448 | 0.039* |
|  | C | 18 | 527 | 8 | 126 | 32 | 939 | 0.278 |
| Dog is lethargic | A | 132 | 9 | 149 | 14 | 1236 | 104 | 0.796 |
|  | B | 89 | 52 | 105 | 58 | 731 | 609 | 0.013* |
|  | C | 7 | 134 | 4 | 159 | 46 | 1294 | 0.465 |
| Dog is confident | A | 1038 | 81 | 260 | 23 | 219 | 21 | 0.630 |
|  | B | 642 | 477 | 153 | 130 | 132 | 108 | 0.529 |
|  | C | 44 | 1075 | 7 | 276 | 6 | 234 | 0.379 |
| Dog seems dull or depressed, not alert | A | 32 | 3 | 83 | 4 | 1403 | 121 | 0.490 |
|  | B | 16 | 19 | 57 | 30 | 853 | 671 | 0.101 |
|  | C | 1 | 34 | 2 | 85 | 55 | 1469 | 0.914 |
| Dog is anxious | A | 502 | 43 | 277 | 22 | 736 | 63 | 0.977 |
|  | B | 308 | 237 | 172 | 127 | 445 | 354 | 0.861 |
|  | C | 12 | 533 | 14 | 285 | 32 | 767 | 0.090 |
| Dog is considered to be very impulsive | A | 421 | 30 | 303 | 27 | 789 | 70 | 0.611 |
|  | B | 242 | 209 | 178 | 152 | 501 | 358 | 0.177 |
|  | C | 14 | 437 | 14 | 316 | 30 | 829 | 0.684 |
| Dog is curious | A | 1361 | 113 | 103 | 11 | 46 | 4 | 0.671 |
|  | B | 822 | 652 | 68 | 46 | 31 | 19 | 0.524 |
|  | C | 54 | 1420 | 2 | 112 | 2 | 48 | 0.628 |
| Dog is affectionate | A | 1392 | 121 | 91 | 4 | 33 | 2 | 0.405 |
|  | B | 846 | 667 | 56 | 39 | 23 | 12 | 0.454 |
|  | C | 56 | 1457 | 1 | 94 | 1 | 34 | 0.411 |
| Dog tends to be calm | A | 890 | 76 | 291 | 29 | 334 | 22 | 0.365 |
|  | B | 562 | 404 | 172 | 148 | 192 | 164 | 0.216 |
|  | C | 32 | 934 | 7 | 313 | 19 | 337 | 0.086 |
| Dog is aloof | A | 251 | 24 | 218 | 15 | 1045 | 89 | 0.636 |
|  | B | 162 | 113 | 134 | 99 | 629 | 505 | 0.550 |
|  | C | 6 | 269 | 7 | 226 | 45 | 1089 | 0.351 |
| Dog has more good days than bad days | A | 1377 | 117 | 47 | 1 | 34 | 4 | 0.264 |
|  | B | 840 | 654 | 27 | 21 | 26 | 12 | 0.344 |
|  | C | 53 | 1441 | 1 | 47 | 3 | 35 | 0.233 |

**Table 10.**
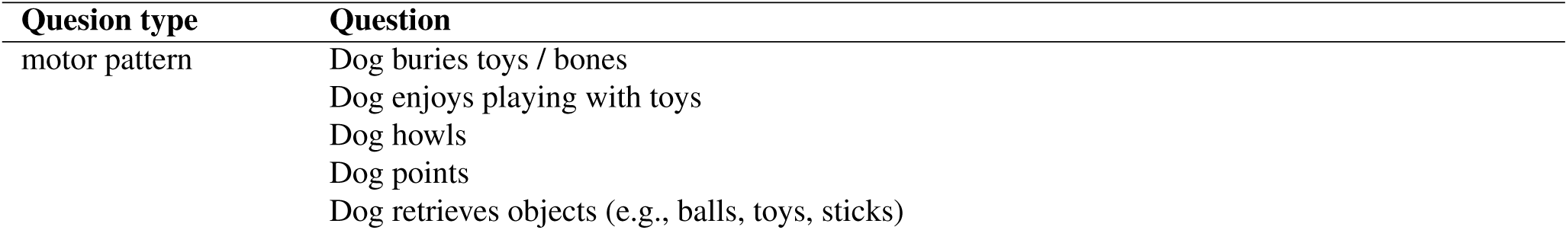
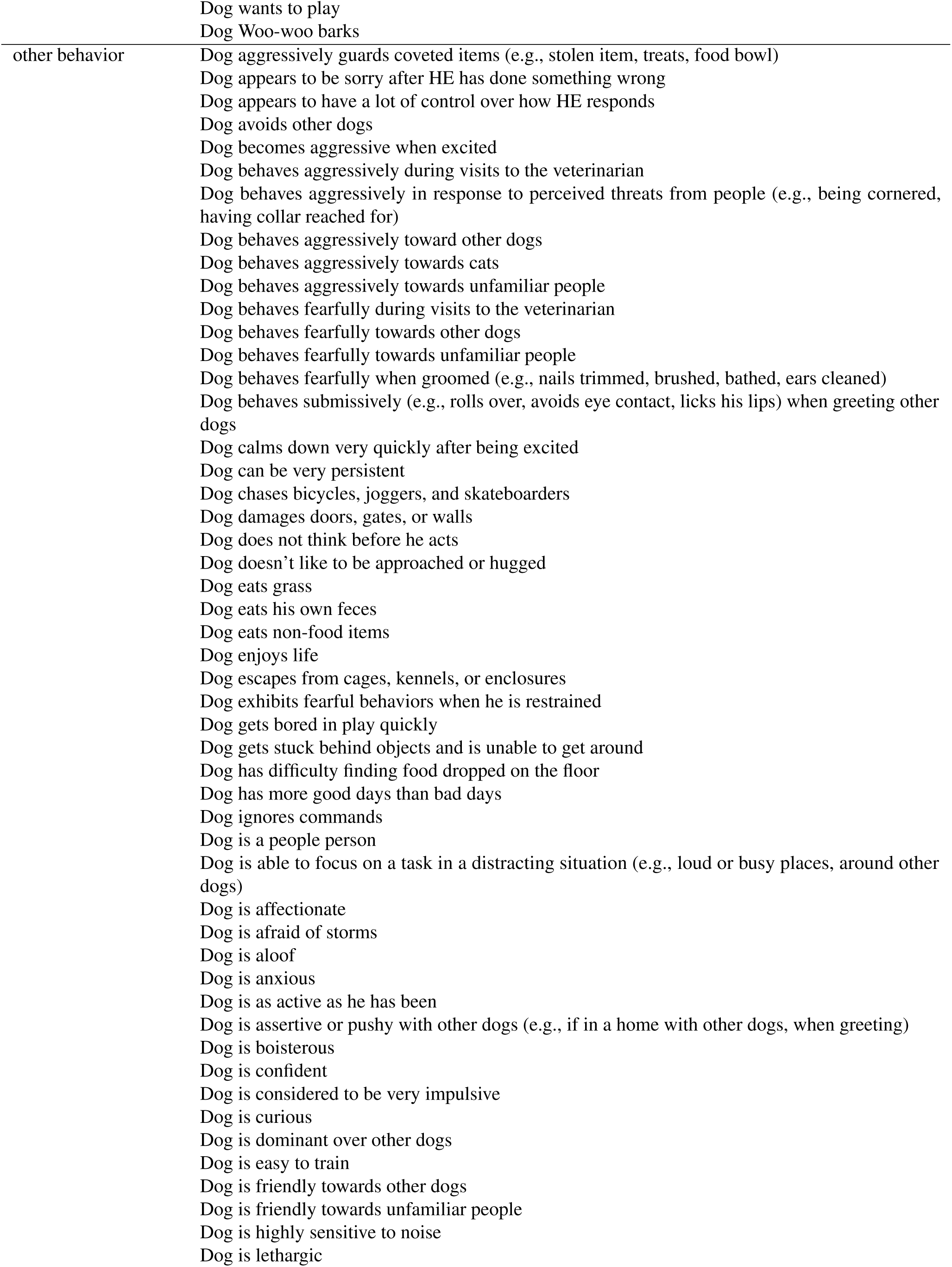
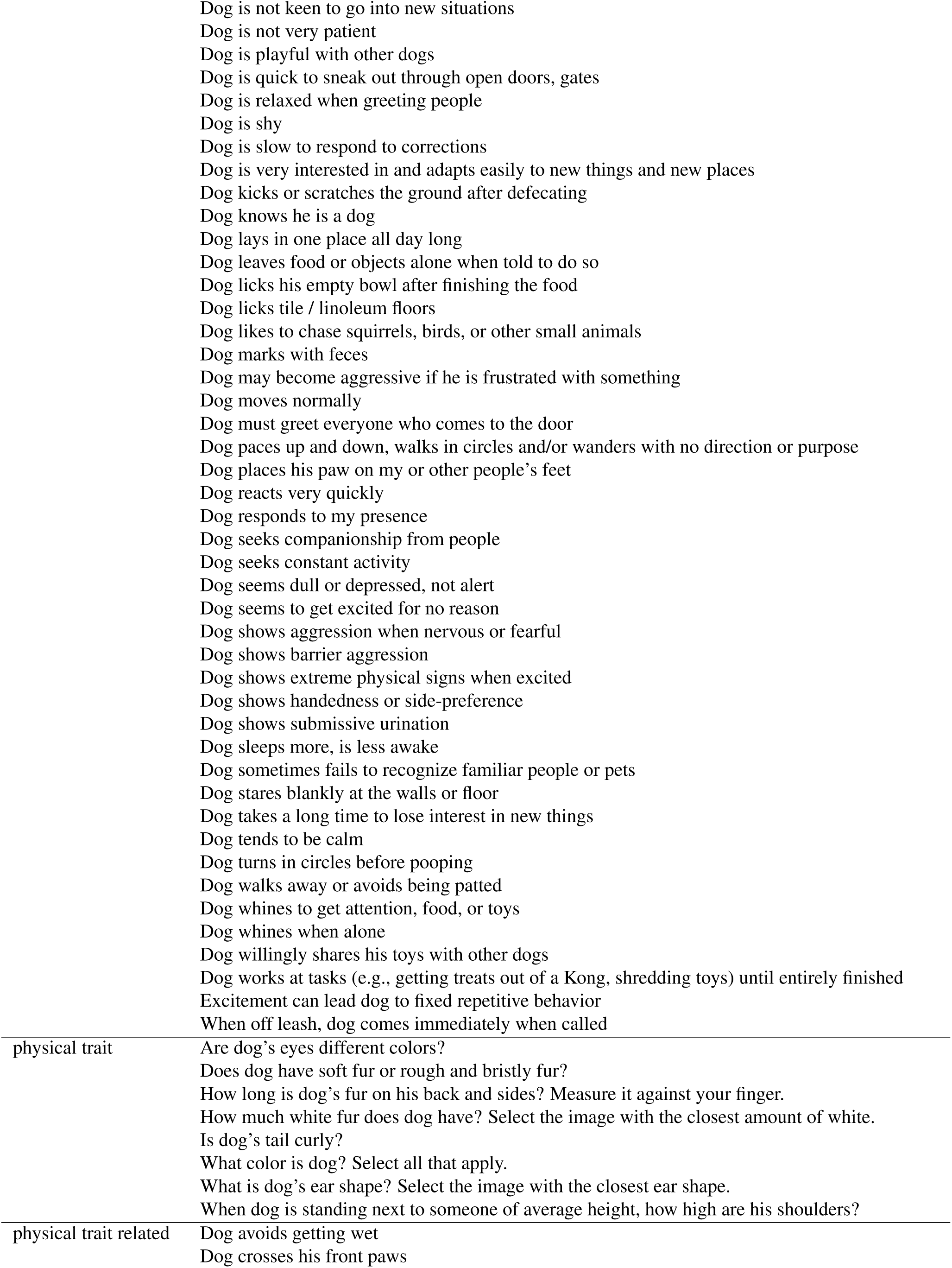
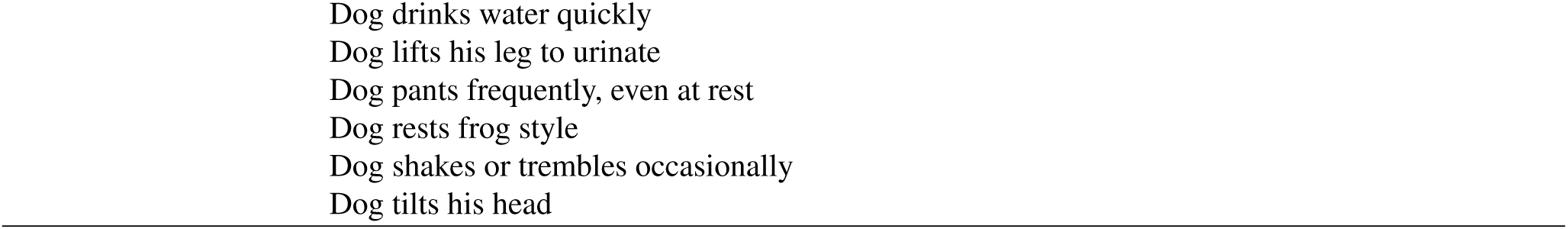
Questions by question types from the original survey.^14^.

**Figure 6.**
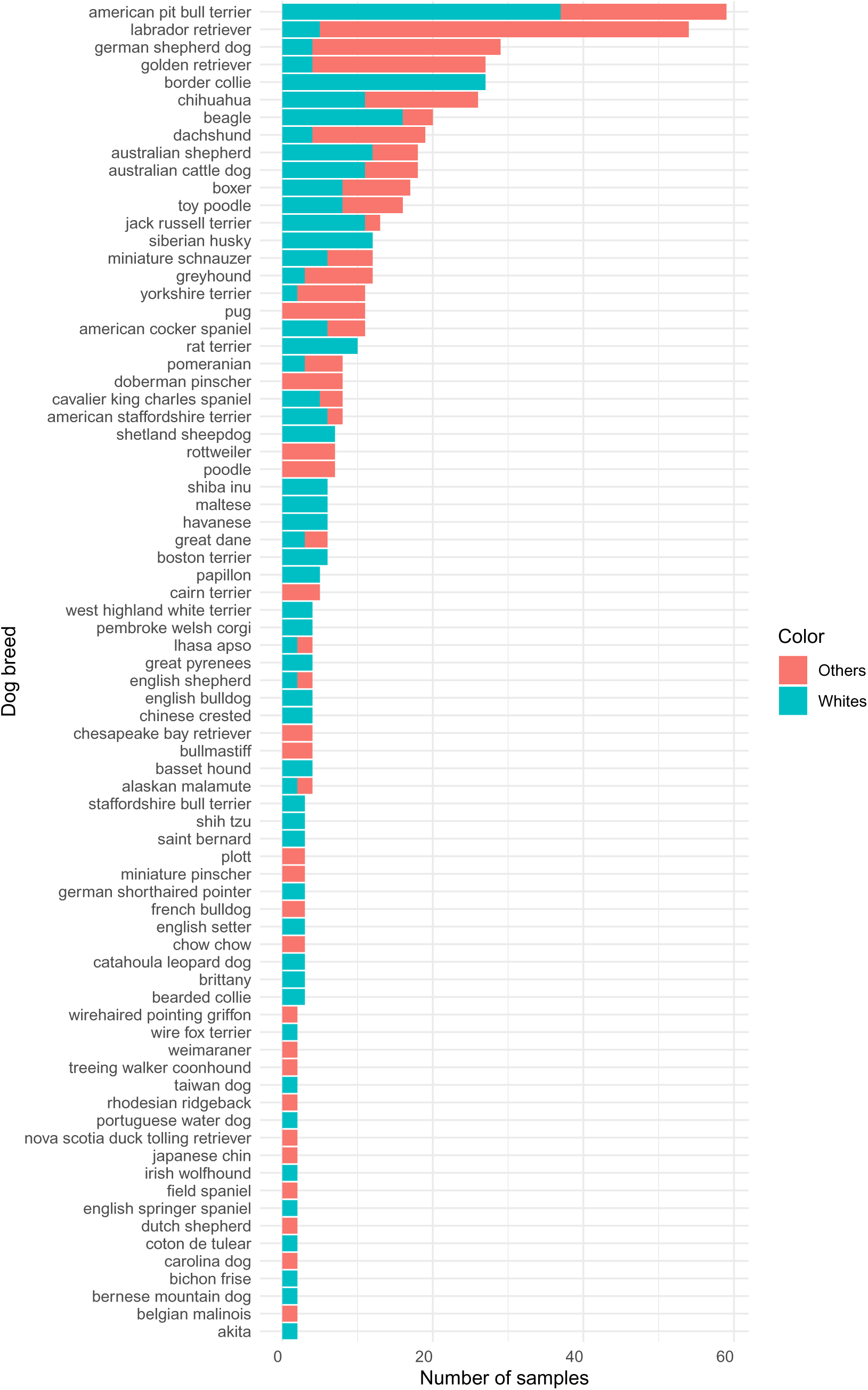
Dog breeds answered to be white (blue) and other color (red) and the number of samples they appeared at. Only breeds appearing in more than 1 sample are presented.

**Figure 7.**
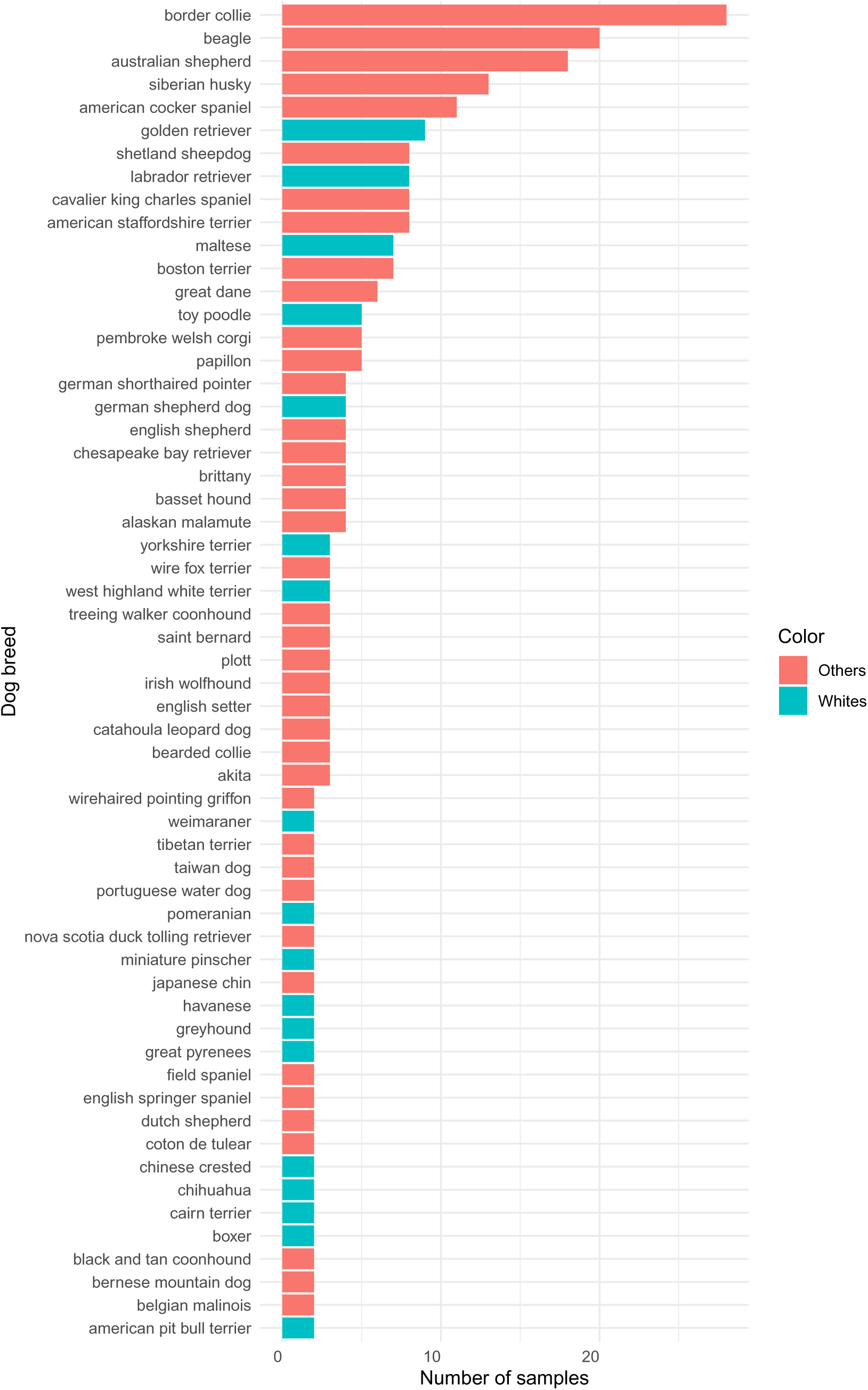
Dog breeds answered to be only white (blue) and not only white (red) and the number of samples they appeared at. Only breeds appearing in more than 1 sample are presented.

